# Rv3723/LucA coordinates fatty acid and cholesterol uptake in *Mycobacterium tuberculosis*

**DOI:** 10.1101/121780

**Authors:** Evgeniya V. Nazarova, Christine R. Montague, Thuy La, Kaley M. Wilburn, Neelima Sukumar, Wonsik Lee, Shannon Caldwell, David G. Russell, Brian C. VanderVen

## Abstract

Pathogenic bacteria have evolved highly specialized systems to extract essential nutrients from their hosts and *Mycobacterium tuberculosis* (Mtb) scavenges lipids (cholesterol and fatty acids) to maintain infection in mammals. While the uptake of cholesterol by Mtb is mediated by the Mce4 transporter, the route(s) of uptake of fatty acids remain unknown. Here, we demonstrate that an uncharacterized protein LucA, integrates the assimilation of both cholesterol and fatty acids in Mtb. LucA interacts with subunits of the Mce1 and Mce4 complexes to coordinate the activities of these nutrient transporters. We also demonstrate that Mce1 functions as an important fatty acid transporter in Mtb and we determine that the integration of cholesterol and fatty acid transport by LucA is required for full bacterial virulence *in vivo*. These data establish that fatty acid and cholesterol assimilation are inexorably linked in Mtb and reveals a key role for LucA in coordinating both transport activities.

## Introduction

*Mycobacterium tuberculosis* (Mtb), the causative agent of human tuberculosis (TB), is responsible for more than 1 million deaths each year and currently infects nearly ∼1.5 billion individuals. A hallmark of TB is that infected individuals rarely develop active TB disease and most infections (∼90%) remain asymptomatic or latent. Mtb is exquisitely adapted to the human host and the bacterium’s ability to metabolize host-derived lipids (fatty acids and cholesterol) aids in bacterial survival and persistence. Therefore elucidating the mechanisms involved in nutrient uptake and metabolism in Mtb will help us better understand the host-pathogen interactions and may identify new vulnerabilities for drug discovery.

Multiple lines of evidence indicate that host lipids (fatty acids and cholesterol) serve as critical carbon sources for Mtb during infection. Mtb propagated in mammalian tissues preferentially metabolizes fatty acids (Bloch and Segal, 1956) and studies using bacterial mutants have repeatedly demonstrated that lipid metabolism promotes Mtb survival during infection (Marrero et al., 2010; McKinney et al., 2000; Muñoz-Elías et al., 2006). Mtb also has the ability to metabolize cholesterol (Van der Geize et al., 2007) and utilization of this nutrient is critical for Mtb survival within macrophages (MΦs) (VanderVen et al., 2015) and in different *in vivo* infection models (Chang et al., 2007; Hu et al., 2010; Nesbitt et al., 2010; Pandey and Sassetti, 2008; Yam et al., 2009).

Mtb’s remarkable capacity to assimilate and metabolize fatty acids is considered a defining characteristic of this pathogen. However, the mechanisms underlying fatty acid uptake in Mtb have remained undetermined. The mycolic acid-containing cell envelope of Mtb constitutes a unique barrier for the import of hydrophobic molecules and likely explains why the Mtb genome does not encode canonical lipid transporters typically found in other bacterial systems (Black et al., 1987; Theodoulou et al., 2016; van den Berg et al., 2004). Mtb imports cholesterol across the mycobacterial cell envelope via the multi-subunit transporter termed Mce4 (Pandey and Sassetti, 2008), and the Mtb genome contains four total unlinked *mce* loci (*mce1-mce4*). While the functions of proteins encoded in the *mce1-mce3* loci are unknown; the similarities shared across the *mce1-4* loci suggest that these loci all encode transporters responsible for the assimilation of hydrophobic molecules.

Most of the genes required for Mtb growth on cholesterol as a sole carbon source have been mapped (Griffin et al., 2011) but, little is known about how Mtb utilizes cholesterol in the presence of other nutrients. Mtb can co-metabolize simple carbon substrates *in vitro* (de Carvalho et al., 2010) suggesting that the bacterium also may utilize fatty acids and cholesterol simultaneously and the bacterium likely encounters both of these substrates together *in vivo*. For example, during infection Mtb induces the formation of foamy MΦs (Peyron et al., 2008) and this process is thought to drive the accumulation of both fatty acids and cholesterol within Mtb infected lesions and MΦs (Kim et al., 2010). Additionally, during infection Mtb uses fatty acids to balance the assimilation of cholesterol-derived intermediates (Lee et al., 2013) suggesting that cholesterol and fatty acid utilization are integrated processes.

Here we performed a forward genetic screen to identify Mtb mutants defective in cholesterol utilization in the presence of fatty acids. We identified LucA, a protein of unknown function and determined that this protein is required for the intake of both fatty acids and cholesterol by Mtb during infection and in axenic culture. We renamed Rv3723 as LucA (<u>l</u>ipid <u>u</u>ptake <u>c</u>oordinator <u>A</u>). LucA facilitates fatty acid and cholesterol uptake in Mtb by stabilizing subunits of the Mce1 and Mce4 transporters. We also determined that the Mce1 complex transports fatty acids in Mtb and that LucA is required for full virulence *in vivo*. Together, these findings demonstrate that fatty acid and cholesterol import in Mtb is integrated via LucA and that coordination of these activities is necessary to support Mtb pathogenesis.

## Results

### Identifying cholesterol utilization genes in Mtb

Using a forward genetic screen we identified genes involved in cholesterol utilization when Mtb is grown in media containing a mixture of fatty acids and carbohydrates. Mtb lacking Icl1 (Mtb Δ*icl1*) fails to grow in rich media containing cholesterol. This growth inhibition is linked to the accumulation of one or more cholesterol-derived intermediates that accumulate when Icl1 is nonfunctional (Eoh and Rhee, 2014; Lee et al., 2013; VanderVen et al., 2015). We reasoned that mutations in the cholesterol utilization pathway would rescue the cholesterol-dependent growth inhibition of Δ*icl1* Mtb (Figure S1A). Therefore, we isolated transposon rescue mutants in a Δ*icl1* Mtb background that gained the ability to grow on media containing cholesterol. In total, 133 clones were isolated and mutations mapped to 19 separate genes (Table S1). The rescue phenotype was confirmed for 16 of the 19 mutants in liquid media containing cholesterol (Figure S1B).

9 of the 16 genes identified had been predicted to be required for growth of Mtb on cholesterol as a sole carbon source (Griffin et al., 2011). These include *rv1130/prpD* and *rv1131/prpC* (Table S1) that enable propionyl-CoA flux into the MCC and thus limits production of toxic MCC intermediates that accumulate in the absence of Icl1 (Eoh and Rhee, 2014; VanderVen et al., 2015). This screen also identified mutations in genes encoding cholesterol catabolic enzymes *rv3545c/cyp125, rv3568c/hsaC*, and *rv3570c/hsaA* (Capyk et al., 2009; Dresen et al., 2010; Yam et al., 2009). These mutations likely rescue growth by reducing the amount of propionyl-CoA released from cholesterol and are consistent with the observation that chemically inhibiting the PrpC or the HsaAB enzymes is sufficient to rescue growth of the Δ*icl1* Mtb in cholesterol (VanderVen et al., 2015).

### *lucA* encodes a membrane protein involved in cholesterol metabolism

17 sibling mutants carrying transposon insertions in the *lucA* gene (Δ*icl1* Tn::*lucA*) were identified with the screen (Table S1). Orthologs of *lucA* are restricted to *Mycobacterium* spp., and this gene encodes an uncharacterized putative integral membrane protein with four predicted transmembrane domains. Previous genetic epistasis studies implicated LucA in cholesterol utilization *in vivo* (Joshi et al., 2006). We confirmed that the transposon insertion in the *lucA* gene is responsible for the growth rescue in the Δ*icl1* background by complementation (Figure 1A). To ascertain LucA’s subcellular localization LucA was fused to green fluorescent protein (LucA-GFP) and expressed in a wild type strain of Mtb that constitutively expresses mCherry. Confocal microscopy revealed a peripheral distribution for LucA-GFP, is consistent with a cell membrane or cell envelope localization (Figure S2).

**Figure. 1.**
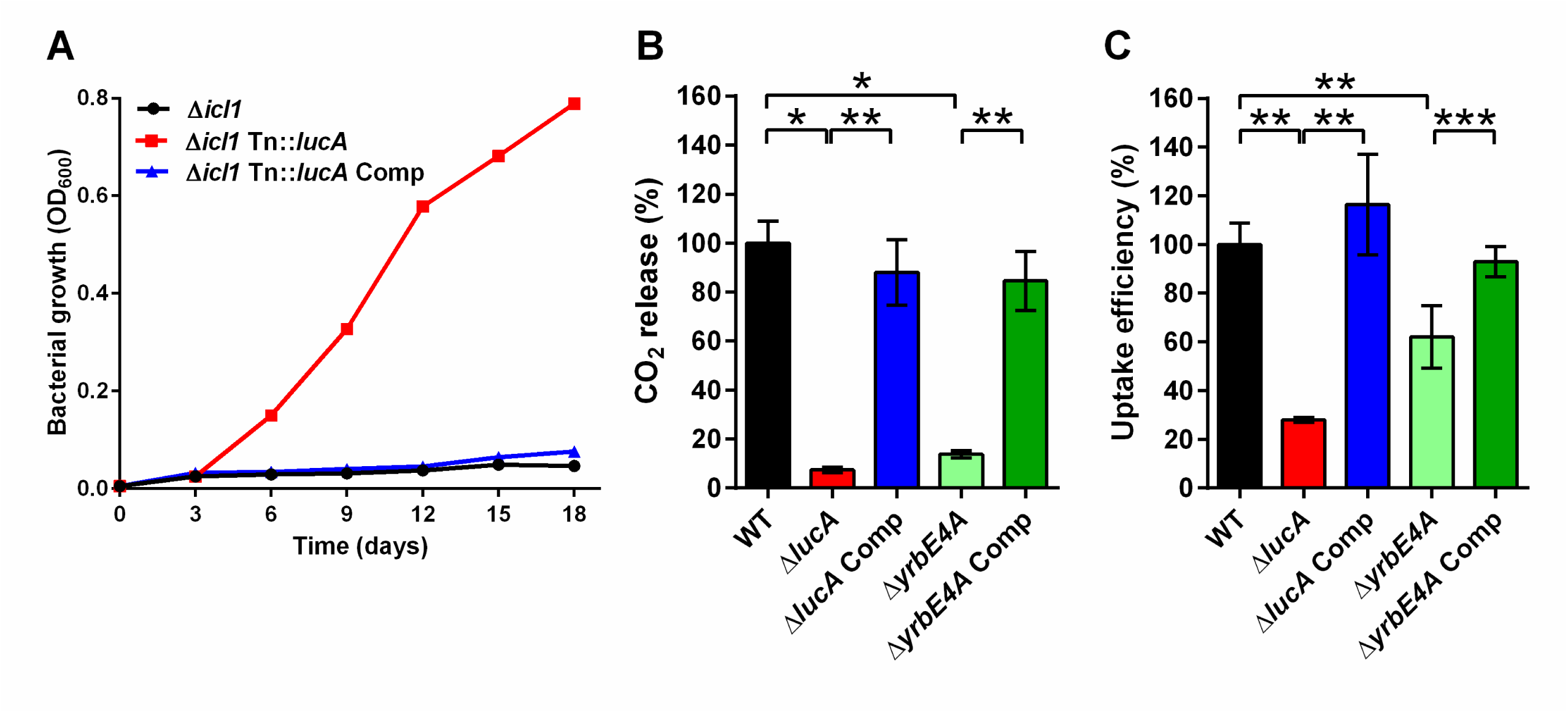
LucA facilitates cholesterol uptake in Mtb. (A) Expressing wild type *lucA* restores the cholesterol-dependent toxicity phenotype and prevents growth of the Δ*icl1* Tn::*lucA* mutant in cholesterol media. (B) The Δ*lucA* and Δ*yrbE4A* mutants are defective in cholesterol metabolism. The catabolic release of ^14^CO_2_ from [4-^14^C]-cholesterol was quantified over 5 hours for each strain. The amount of ^14^CO_2_ released by wild type was set to 100% and the amount of ^14^CO_2_ released by the other strains was expressed as a ratio relative to wild type. (C) The Δ*lucA* and *ΔyrbE4A* mutants are defective in cholesterol uptake. Mtb associated cholesterol was quantified using [4-^14^C]-cholesterol over 2 hours. The rate of uptake was determined using linear regression (Figure S5 and Table S2). The cholesterol uptake rate in wild type was set to 100% and the uptake rates for the other strains were expressed as a ratio relative to wild type. (B and C) Data are means ± SD (n = 4). *p < 0.0005, **p < 0.005, ***p < 0.05 (Student’s t test).

### LucA facilitates cholesterol uptake and metabolism

Given the potential role of LucA in cholesterol metabolism, we constructed LucA mutant in Mtb Erdman by allelic exchange for further analysis (Mtb Δ*lucA*) (Figure S3) and confirmed that *lucA* is required for the catabolic release of ^14^CO_2_ from [4-^14^C]-cholesterol using radiorespirometry (VanderVen et al., 2015). This assay detected a 90% reduction in the amount of ^14^CO_2_ released from Mtb Δ*lucA* relative to the wild type and the complemented strains (Figure 1B). The reduction in ^14^CO_2_ release was not due to a growth defect (Figure S4) and the percent ^14^CO_2_ released was normalized to bacterial biomass.

To determine the impact of LucA on cholesterol uptake in Mtb we quantified the rate of [4-^14^C]-cholesterol uptake using an established assay (Pandey and Sassetti, 2008). We found that the Δ*lucA* mutant assimilates 70% less cholesterol relative to the wild type and complemented controls (Figure 1C, Table S2, and Figure S5). The level of cholesterol uptake observed with Δ*lucA* mutant was comparable to the amount of cholesterol assimilated by a Mce4 mutant (Δ*yrbE4A*) (Figure 1C and Table S2).

### The *ΔlucA* mutant is defective in cholesterol utilization during infection

Analysis of the bacterial transcriptional responses during infection revealed a role for LucA in cholesterol utilization. Expression of KstR regulon genes were strongly down-regulated in the Δ*lucA* mutant in comparison to the wild type and complemented strains (Figure 2, Table S3, and Table S4). KstR is a repressor that controls the expression of genes encoding enzymes that metabolize the side chain and A/B rings of cholesterol (Kendall et al., 2007). The KstR regulon is activated in a “feed forward” manner when KstR is de-repressed by binding to the early cholesterol degradation intermediate, 3-oxocholest-4-en-26-oyl-CoA (Ho et al., 2016) during infection (Rohde et al., 2012; Schnappinger et al., 2003). Thus, the down-regulation of the KstR regulon in the Δ*lucA* mutant indicates a decrease in cholesterol uptake.

**Figure. 2.**
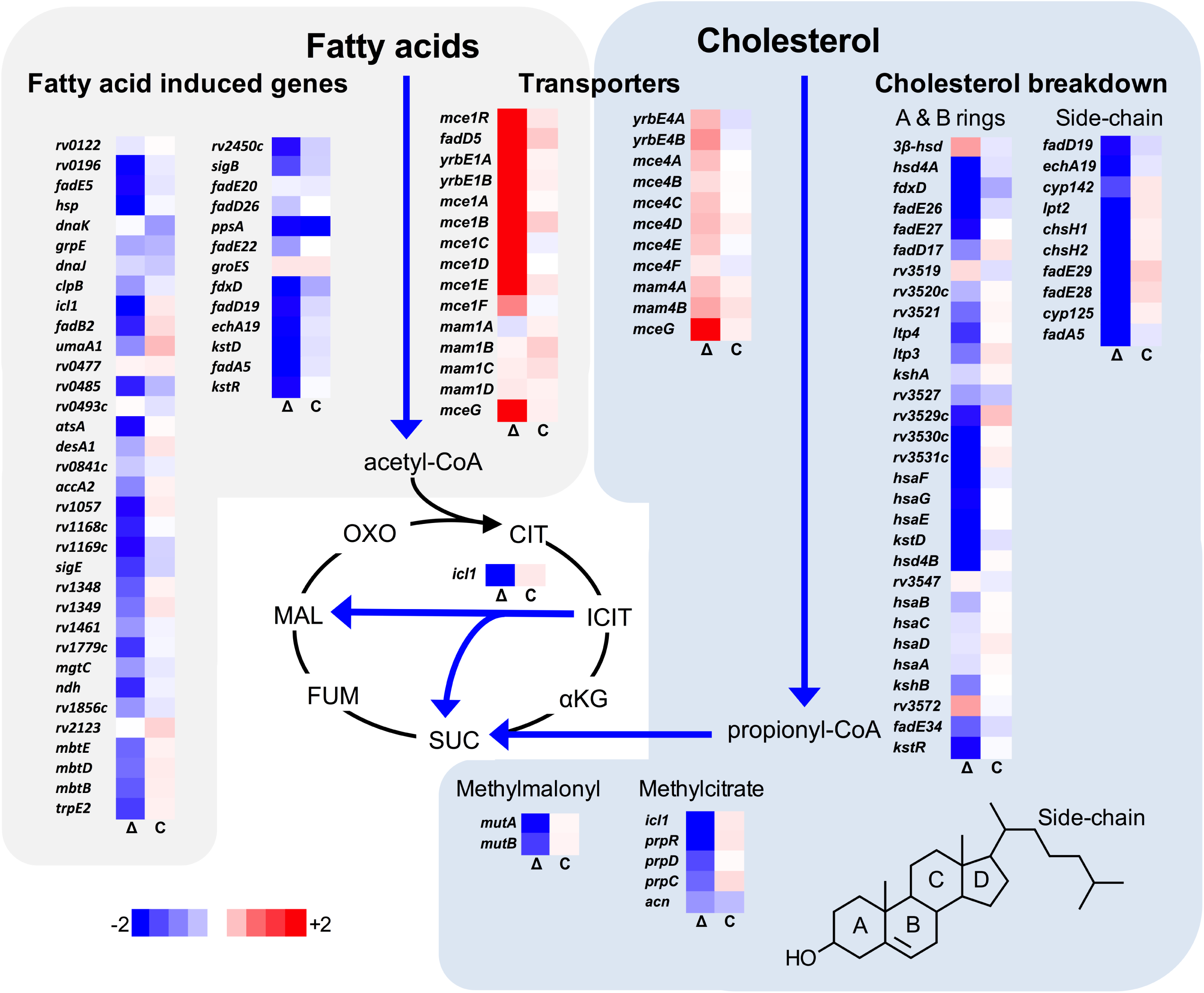
Transcriptional profile of the Δ*lucA* mutant during infection in MΦs. Bacterial gene expression profiles were determined at day 3-post infection in resting murine bone marrow-derived MΦs. Gene expression values from the Δ*lucA* mutant (Δ) and complement (C) strains were normalized to wild type. Genes in the KstR regulon are described (Kendall et al., 2007), and the fatty acid induced genes are up-regulated >2.5-fold in response to palmitate and >1.4-fold in MΦs (Schnappinger et al., 2003). Data are means (n = 3) and the absolute values and statistical analysis are shown in (Tables S3, S4, and S5).

Mtb assimilates cholesterol-derived propionyl-CoA into central metabolism via the methyl-malonyl pathway (MMP) and the MCC and these genes are highly expressed when Mtb metabolizes cholesterol (Griffin et al., 2012; Savvi et al., 2008). We found that expression of the MMP and MCC genes were also down-regulated in the Mtb Δ*lucA* mutant (Figure 2). Cholesterol degradation in Mtb produces propionyl-CoA, which activates the *rv1130/prpD* promoter (Griffin et al., 2012; Masiewicz et al., 2012). To monitor cholesterol breakdown in Mtb during infection we constructed a reporter plasmid where the *rv1130/prpD* promoter controls GFP expression and mCherry is constitutively expressed (*prpD*’::GFP smyc’::mCherry). We validated this reporter by determining that expression of GFP was induced >20-fold in wild type Mtb when the bacteria were grown in media containing cholesterol or propionate (Figure S6A and Figure S6B). Using this reporter we observed a ∼75% reduction in GFP expression from the Δ*lucA* mutant relative to the wild type and complement controls when the bacteria were grown in media containing cholesterol (Figure S6C). We also quantified GFP expression in the Mtb Δ*lucA* mutant during infection in MΦs. Intracellular bacteria were isolated from the infected MΦs and bacterial GFP expression levels were determined by flow cytometery. This analysis demonstrated a ∼75% reduction in the GFP signal from the Mtb Δ*lucA* mutant (Figure S6D) supporting the idea that the Mtb Δ*lucA* mutant is defective in utilizing cholesterol during infection.

### The Mtb *ΔlucA* mutant does not assimilate fatty acids during infection

Unexpectedly, the transcriptional response in the Mtb Δ*lucA* mutant also revealed a gene expression signature consistent with a defect in fatty acid utilization (Table S3 and Table S5). For this, we focused on the Mtb genes induced in the bacteria during MΦ infection and by palmitate, and observed that the majority of these genes were strongly down-regulated in the Δ*lucA* mutant (Figure 2).

To verify the fatty acid utilization defect in the Δ*lucA* mutant we quantified the uptake of fluorescent palmitate (Bodipy-C16) by the intracellular bacteria. Resting MΦs were infected with wild type, Δ*lucA* mutant, and complemented strains, all constitutively expressing mCherry. On day 3 of infection the MΦs were pulsed with Bodipy-C16. Confocal analysis revealed that the wild type and the complemented bacteria accumulated intracellular Bodipy-C16 as visible punctate inclusions, while the Δ*lucA* mutant did not (Figure 3A). To corroborate this finding, intracellular bacteria were isolated from pulse-labeled MΦs and Bodipy-C16 assimilation by Mtb was quantified by flow-cytometery. This analysis revealed a 10-fold reduction in the amount of Bodipy-C16 assimilated by the Mtb Δ*lucA* mutant relative to the wild type and complemented strains (Figure 3B). There was minimal difference in bacterial colony forming units (CFU) at day 3 of these experiments indicating that the decreased levels of assimilated Bodipy-C16 were not due to a loss of bacterial viability (Figure S7).

**Figure 3.**
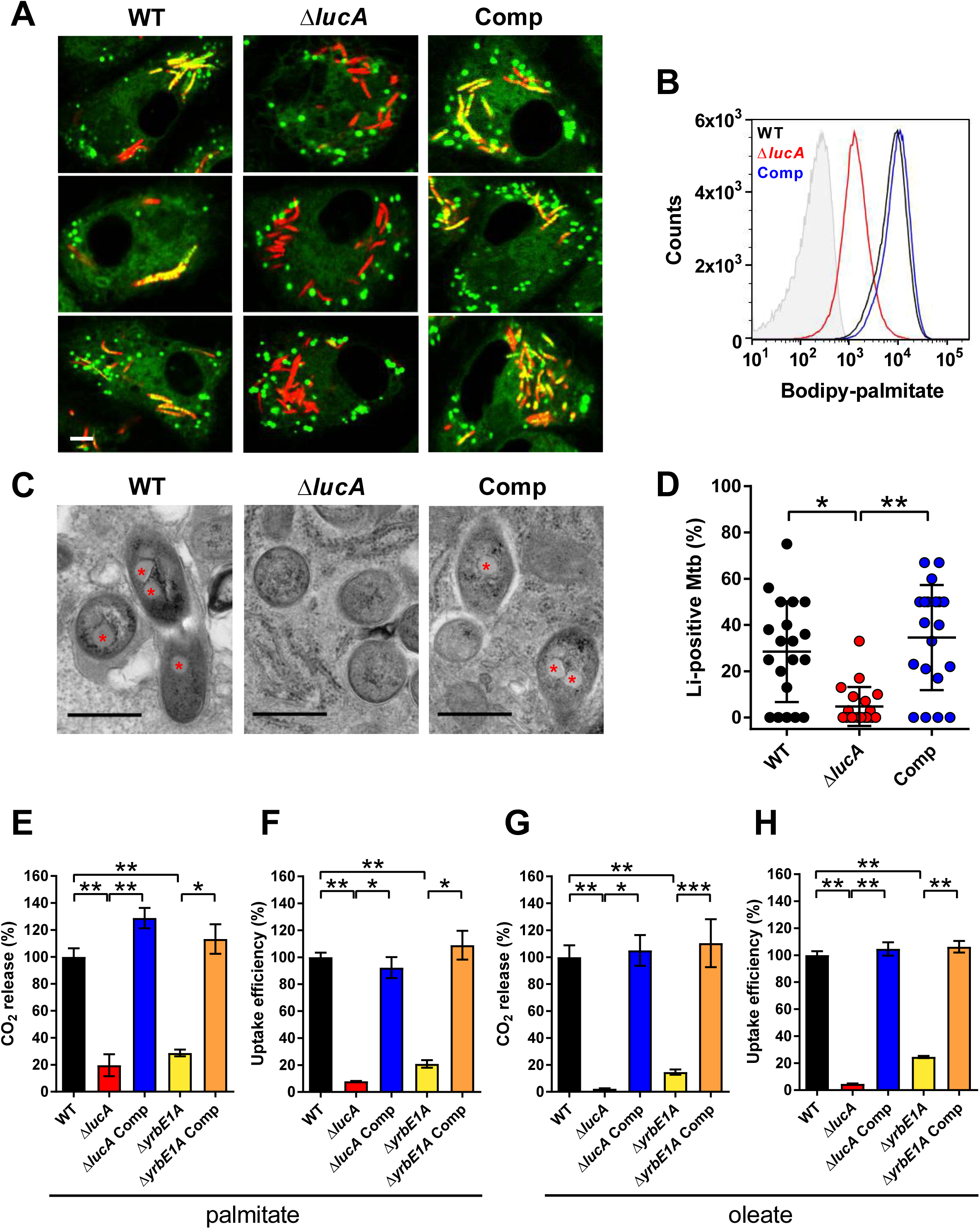
LucA facilitates fatty acid uptake during infection and in axenic culture. (A) The Δ*lucA* mutant does not accumulate Bodipy-C16 in cytosolic lipid inclusions compared to wild type and the complemented strain. Representative confocal images of infected resting murine MΦs pulse labeled with Bodipy-C16 (red = mCherry Mtb, green = Bodipy-C16). Scale bar 5.0 μm. (B) Flow cytometery based quantification of Bodipy-C16 incorporation by Mtb isolated from pulse labeled MΦs. Shaded histogram represents the Mtb auto-fluorescence in the green channel. 100,000 events from each strain were analyzed. (C) Transmission electron microscopy reveals a lack of intracellular Li’s in the Δ*lucA* mutant at day 3-post infection in MΦs. Li’s indicated by asterisks are readily apparent in wild type and the complemented strain but not in the Δ*lucA* mutant. Scale bar 0.5 μm. (D) Quantification of intracellular Mtb containing visible Li’s per macrophage section. Horizontal bars are means ± SD (n = 20). (E, G) The Δ*lucA* and Δ*yrbE1A* mutants are defective in fatty acid metabolism. The catabolic release of ^14^CO_2_ from [^14^C(U)]-palmitic acid (E) or [1-^14^C]-oleic acid (G) was quantified over 5 hours for each strain. (F, H) The Δ*lucA* and Δ*yrbE1A* mutants are defective in fatty acid uptake. Mtb associated fatty acids were quantified using [^14^C(U)]-palmitic acid (F) or [1-^14^C]-oleic acid (H) over 2 hours for each strain. (E-H) Data was calculated as in Figure 1B and C. Data are means ± SD (n ≥ 4). *p < 0.0005, **p < 0.0001, ***p < 0.005 (Student’s t test).

It is possible that the Δ*lucA* mutant traffics to lysosomal compartments in the MΦ, which could potentially restrict access to Bodipy-C16. To rule this out lysosomes were pulse labeled with Alexa 647-dextran prior to infection. At day 3 of infection we determined that the Δ*lucA* mutant did not colocalize with the Alexa 647-dextran-loaded lysosomes with Pearson coefficient of correlation values of 0.052±0.062 (wild type), 0.101±0.054 (Δ*lucA*), and 0.190±0.174 (complement) (Figure S8).

During infection Mtb can sequester fatty acids as triacylglycerol (TAG) within cytosolic intracellular lipid inclusions (Li’s) (Daniel et al., 2011). Therefore, the presence of Li’s can serve as an indicator of fatty acid assimilation by Mtb during infection. To image intracellular Li’s in Mtb we used the neutral lipid stain Bodipy-493/503 (Listenberger and Brown, 2007). Staining infected murine MΦs with Bodipy-493/503 revealed a punctate staining pattern inside the wild type and complemented strains while no staining was observed in the Mtb Δ*lucA* mutant (Figure S9A). Additionally, Li’s are also visible by transmission electron microscopy (Figure 3C) and only ∼5% of the intracellular Δ*lucA* mutant cells had identifiable Li’s, while 25% of the wild type and 35% of complemented bacteria had visible Li’s during infection (Figure 3D).

The decrease in Li’s within the Δ*lucA* mutant could also be due to enhanced turnover of the intracellular Li’s. To rule this possibility out, intracellular bacteria were treated with the broad-spectrum lipase inhibitor, tetrahydrolipstatin (THL), which can inhibit TAG turnover in Mtb (Baek et al., 2011). We predicted that if TAG were more efficiently degraded in the Mtb Δ*lucA* mutant, THL treatment would restore the presence of intracellular Li’s in the mutant. There was no increase in the levels of intracellular Li’s in the Mtb Δ*lucA* mutant following THL treatment (Figure S9B). These observations indicate that the Mtb Δ*lucA* mutant is unable to utilize both fatty acids and cholesterol during infection in MΦs.

### LucA facilitates fatty acid uptake

We next quantified fatty acid metabolism and uptake in the Mtb Δ*lucA* mutant using radiolabeled fatty acids. Fatty acid metabolism was measured by quantifying the oxidative release of ^14^C-CO_2_ from [^14^C(U)]-palmitate and [1-^14^C]-oleate by radiorespirometry. This assay detected a ∼80% and ∼95% reduction in the Δ*lucA* mutant’s ability to metabolize the [^14^C(U)]-palmitate and [1-^14^C]-oleate, respectively (Figure 3E and 3G).

Fatty acid uptake was quantified with [^14^C(U)]-palmitate and [1-^14^C]-oleate as described (Forrellad et al., 2014). Relative to the wild type and complemented strains we observed a ∼90% and ∼95% reduction in the Δ*lucA* mutant’s ability to assimilate [^14^C(U)]-palmitate and [1-^14^C]-oleate, respectively (Figure 3F, Figure 3H, and Figure S2). The ^14^C-fatty uptake rate was calculated (Figure S5) and normalized to biomass. Based on these results we conclude that deletion of LucA perturbs fatty acid uptake by the bacterium, which decreases the bacterium’s ability to metabolize fatty acids.

### Mce1 is a fatty acid transporter

The system(s) responsible for fatty acid import in Mtb have remained elusive. While the Δ*lucA* mutant is defective in fatty acid uptake, the LucA protein lacks any recognizable domains that would predict a transport function for the protein. We hypothesized that, in the absence of LucA, Mtb may up-regulate expression of an actual fatty acid transporter in attempt to compensate for the fatty acid uptake defect. Upon analysis, we observed that genes in the *mce1* loci are strongly induced in Δ*lucA* mutant in MΦs (Figure 2). To test the hypothesis that Mce1 functions as a fatty acid transporter in Mtb, we generated a Mce1 mutant by deleting the Mce1 permease Rv0167/YrbE1A subunit (Δ*yrbE1A*) and quantified fatty acid uptake and metabolism with this strain. The Δ*yrbE1A* mutant displayed a ∼70% and ∼85% reduction in its ability to metabolize [^14^C(U)]-palmitate and [1-^14^C]-oleate, respectively (Figures 3E, 3G, and S2). Relative to the wild type and complement strains we detected a ∼80% and ∼75% reduction in the Δ*yrbE1A* mutant’s ability to intake [^14^C(U)]-palmitate and [1-^14^C]-oleate (Figure 3F and 3H). Notably, Δ*yrbE1A* was able to transport and metabolize cholesterol to wild type levels (Figure S10). Lastly, a mutant lacking 8 genes of the *mce1* loci (Δmce1) was defective in fatty acid uptake and metabolism, but had no detectable defect in cholesterol uptake or metabolism (Figure S11). Based on these results we conclude that the Mce1 complex functions as a dedicated fatty acid transporter in Mtb.

### LucA interacts with Mce1- and Mce4-associated proteins

To shed light on the function of LucA we conducted a mycobacterial 2-hybrid screen to identify protein fragments that interact with LucA (Singh et al., 2006). For this, the F_3_ domain of murine dihydrofolate reductase (mDHFR) was fused to the C-terminus of a full length LucA (LucA-F_3_) (Table S6). We co-expressed LucA-F_3_ in *M. smegmatis* (Msm) along with a library (2×10^6^) of random Mtb protein fragments fused to the F_1,2_ domain of mDHFR. Interactions between bait and prey fusions containing the split domains of mDHFR confer resistance to trimethoprim (TRIM). This screen identified 4 sibling clones which encode an in-frame N-terminal fragment (1-110 aa) of *rv3492c/mam4B* fused to the F_1,2_domain of mDHFR. Mam4B is encoded by a gene within the *mce4* loci (Figure 4A) and this protein is predicted to function as a subunit of the Mce4 cholesterol transport complex. Thus, we re-cloned Mam4B fused to the F_1,2_ domain of mDHFR (Mam4B(TR)-F_1,2_) (1-75 aa) and confirmed that co-expressing Mam4B(TR)-F_1,2_ with LucA-F_3_ in Msm conferred trimethoprim (TRIM) resistance (Figure 4B and Table S7). Co-expressing just the transmembrane domain of Mam4B fused to the F_1,2_ domain of mDHFR (Mam4B(TM)-F_1,2_) (1-70 aa) conferred TRIM resistance (Figure 4B and Table S6) indicative of a biologically-significant interaction. The Mtb genome encodes additional homologs of Rv3492c/Mam4B that are associated with the Mce1 and Mce4 transporters (Figure 4A). To determine if these subunits also interact with LucA we generated F_1,2_ fusions to Rv0177/Mam1C and Rv0199/OmamA and found that co-expression of these along with LucA-F_3_ conferred TRIM resistance (Figure 4B, Table S6). These data indicate that LucA physically interacts with subunits of the Mce1 and Mce4 transporters suggesting that the LucA protein participates in the function of these complexes.

**Figure 4.**
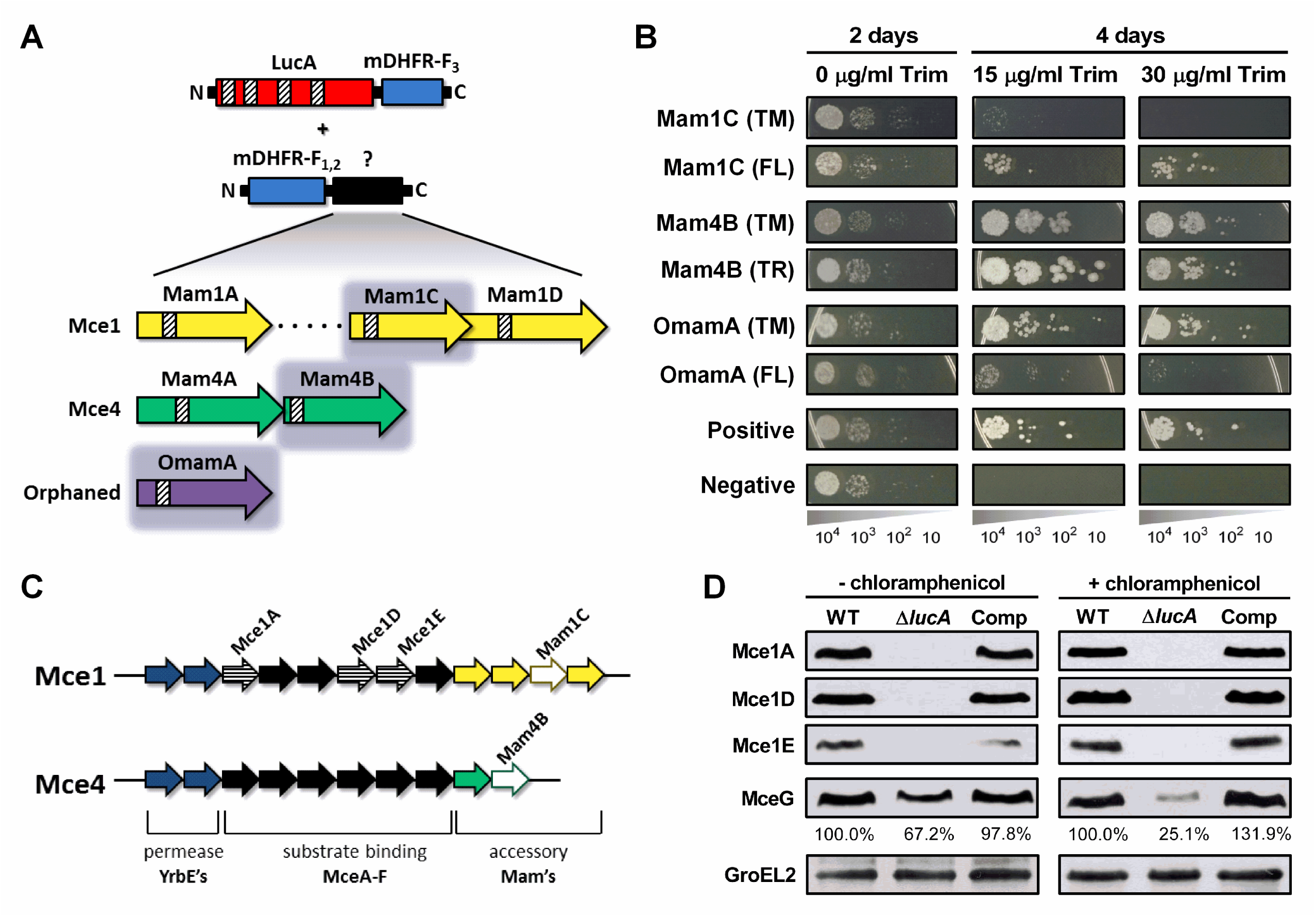
LucA interacts with subunits of Mce1 and Mce4 transporters and is required for their stability. (A) Schematic of proteins tested for interaction with LucA. Top, schematic of protein fusions with DHFR domains. Bottom, putative accessory subunits of Mce transporters homologous to Mam4B. Shading indicates proteins tested positive for interaction with LucA. Striped boxes indicate predicted transmembrane domains. (B) Two-hybrid assay for the detected interactions between LucA and accessory proteins of Mce1 and Mce4. Serial dilutions of Msm co-expressing the LucA-F_3_ bait and the indicated prey were spotted onto an agar plate without antibiotic or onto plates containing trimethoprim at the indicated concentration. Growth on trimethoprim plates is indicative of protein-protein interaction. Details prey proteins see (Table S6). Positive: Msm co-expressing *Saccharomyces cerevisiae* homodimeric leucine zipper subunits (GCN4-F_1,2_ and GCN4-F_3_). Negative: Msm expressing LucA-F_3_ and GCN4-F_1,2_. (C) Genome organization of Mce1 and Mce4 subunits. Striped arrows indicate genes encoding proteins tested for stability in Δ*lucA*. Empty arrows indicate genes encoding accessory proteins that interact with LucA. (D) Subunits of the Mce1 and Mce4 complexes are degraded in the Δ*lucA* mutant. Whole cell lysates were prepared from wild type, Δ*lucA*, and the complement strain. The indicated proteins were analyzed separately by SDS-PAGE and western blotting. Chloramphenicol was added for 2 days before protein extraction where indicated. The levels of MceG were quantified and expressed as a ratio relative to levels of MceG in the wild type lysates. GroEL2 is used as a loading control and blots are representative of two independent experiments.

### LucA stabilizes subunits of the Mce1 and Mce4 complex

Recently, it was demonstrated that Rv0199/OmamA is required for cholesterol metabolism in Mtb (Perkowski et al., 2016). Given that LucA interacts with Rv0199/OmamA and the related homologs Rv3492c/Mam4B and Rv0177/Mam1C we hypothesized that both the Mce1 and Mce4 transporters may also be destabilized in the Δ*lucA* mutant. To test this, Mtb was grown under the conditions used in the lipid uptake experiments. Gene expression analysis by qPCR confirmed that the *mce1* and *mce4* genes are expressed in the Δ*lucA* mutant to equivalent levels relative to the wild type and complemented strains under the assay conditions (Figure S12). In contrast, analysis of whole cell lysates revealed that, at the protein level, the putative subunits of the Mce1 complex (including Mce1A, Mce1D, Mce1E) were completely degraded in the Δ*lucA* mutant (Figure 4C and Figure 4D). Unfortunately, thus far we have been unable to raise antibodies specific to the analogous Mce4 subunits to determine if they to are degraded.

It is thought that MceG/Rv0655 functions as a common ATPase to hydrolyze ATP and facilitate uptake through the Mce1 and Mce4 transporters (Joshi et al., 2006; Mohn et al., 2008; Pandey and Sassetti, 2008). Given that the stability of MceG requires co-expression of the Mce1 and Mce4 permease components (Joshi et al., 2006) we monitored MceG protein levels and observed a 30% decrease in amount of MceG in the Δ*lucA* mutant (Figure 4D). The rate of synthesis of Mce subunit proteins can exceed the rates of degradation (Perkowski et al., 2016) and this can mask protein instability. We therefore used chloramphenicol to suppress protein synthesis and under this condition the amount of MceG protein decreased by 75% in the Δ*lucA* mutant (Figure 4D). These results demonstrate that, in the absence of LucA, not only subunits of the Mce1 transporter complex, but also shared MceG ATPase, are degraded. These data provide an explanation for the linked defect in both fatty acid and cholesterol uptake observed in the Δ*lucA* mutant.

### Substrate binding and translocation by Mce4 are dissociable events

Our data indicates that the LucA protein interacts with Rv3492c/Mam4B, a putative subunit of the Mce4 complex and we predict that this protein is required for cholesterol import. To test this hypothesis, Rv3492c/Mam4B was deleted (Δ*mam4B*) and we quantified the rate of [4-^14^C]-cholesterol uptake in this mutant. We found that the level of association of [4-^14^C]-cholesterol with the Δ*mam4B* mutant was comparable to wild type bacteria (Figure S13A). However, analysis of cholesterol breakdown in the Δ*mam4B* mutant revealed an 80% reduction in [4-^14^C]-cholesterol metabolism (Figure S13B). Additionally, the Δ*mam4B* had no defect in fatty acid uptake or metabolism (Figures S13C-E and D-F). Together these data indicate that Rv3492c/Mam4B is required to complete the process of Mce4-mediated cholesterol internalization in Mtb.

### LucA contributes to the *in vivo* fitness of Mtb

These data indicate that LucA is involved in the utilization of two critical host-derived lipid substrates. It is known that deletion of one or both of the Mce1 and Mce4 transporters leads to decreased survival of the pathogen in mouse model infection (Joshi et al., 2006) and we predict that both of these transporters would be nonfunctional in the Δ*lucA* mutant. In human MΦs the Δ*lucA* mutant displayed a growth lag that culminates in a 10-fold difference in bacterial counts compared to wild type and complemented strains over a 7-day infection period (Figure 5A). A similar phenotype was observed in the resting murine MΦs where the Δ*lucA* mutant replicated poorly over a 10-day infection period (Figure 5B). In both human and murine MΦs the final CFU counts for the Δ*lucA* mutant remained close to initial inoculum levels. These data are consistent with the hypothesis that Mtb requires LucA to sustain maximal growth on cholesterol and/or fatty acid substrates in MΦs.

**Figure 5.**
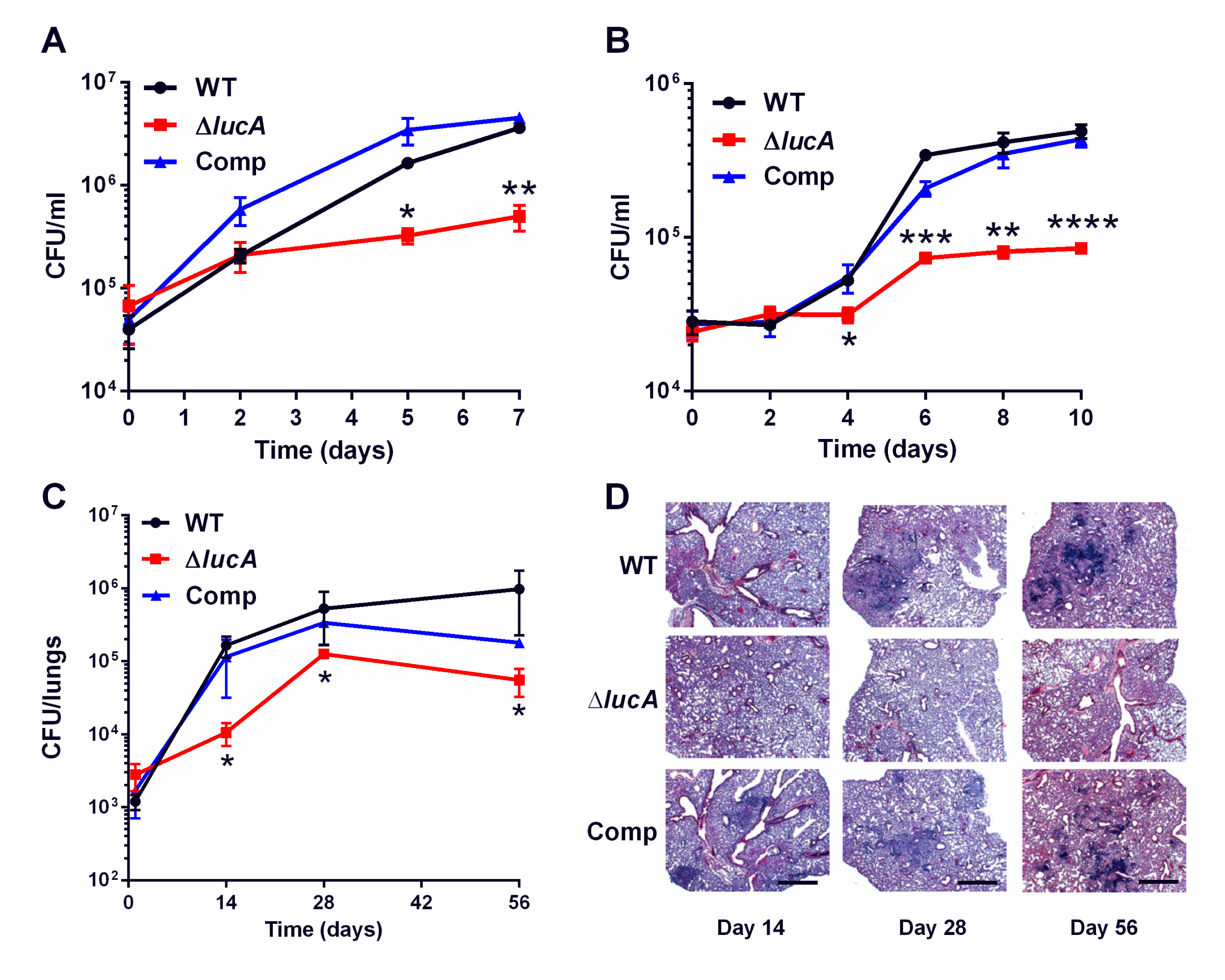
LucA is required for the full fitness of Mtb in MΦs and survival in mouse lungs. (A) Bacterial replication in resting human monocyte derived MΦs. Data are means ± SD (n = 3). (B) Bacterial replication in resting murine bone marrow derived MΦs. Data are means ± SD (n = 3). (C) Bacterial survival in mouse lung tissues. Data are means ± SD (n = 5 per time point) (D) Lung pathology of infected mice collected at indicated time points and H&E stained. Scale bar 400 μm. *p < 0.05, **p < 0.005, ***p < 0.001, ****p < 0.0005 (Student’s t test).

In the lung tissues of C57BL/6J mice, the Δ*lucA* mutant also demonstrated a fitness defect and did not attain levels of bacterial burden comparable to either the wild type or complemented strains. At day 14 post-infection 10-fold fewer CFU’s of the Δ*lucA* mutant were recovered relative to wild type and the complemented strains. During the remainder of the 56-day infection period, growth of the Δ*lucA* mutant remained restricted, leading to 3- to 5-fold reduction in viable Δ*lucA* bacteria (Figure 5C). The Δ*lucA* mutant also induced less pulmonary pathology throughout the entire course of infection (Figure 5D). Complementation almost completely restored pathogenicity and lung tissue pathology confirming the specificity of the mutant phenotype to the *lucA* gene.

## Discussion

While there is general agreement as to the role of Mce4 in cholesterol uptake in Mtb, the process of fatty acid assimilation by the bacterium has remained enigmatic. These data shed new light on the coordination of fatty acid and cholesterol import and reveal that a network of proteins associates with the Mce1 and Mce4 transporters to integrate the uptake of both fatty acids and cholesterol.

Using an unbiased forward genetic screen we discovered that transposon insertions in the *LucA* gene rescue cholesterol toxicity in an Mtb strain that lacks Δ*icl1*. Subsequent analysis confirmed that mutation of *LucA* (Δ*lucA*) has a profound impact on cholesterol uptake (Figure 1B and Figure 1C), and that this cholesterol uptake defect confers growth rescue in the Δ*icl1* Tn::*lucA* double mutant.

Transcriptional profiling revealed that genes normally activated by cholesterol are down-regulated in the Δ*lucA* mutant during infection in MΦs. This is consistent with a defect in cholesterol uptake in this mutant (Figures 2, S3, and S4). We were surprised to discover that the Δ*lucA* mutant also produces a gene expression signature indicative of a fatty acid metabolism defect (Figure 2, S3, and S5). Although the functions for many of these “fatty acid-induced” genes are unknown their expression pattern serves as a reliable indicator of a fatty acid uptake defect in Mtb. The majority of the 20 most highly-expressed genes in the Δ*lucA* during infection in MΦs map to the *mce1* loci (Figure 2, Table S3). Up-regulation of genes encoding the Mce1 fatty acid transporter in the Δ*lucA* mutant during infection likely reflects an attempt by the bacterium to compensate for the absence of fatty acids that normally fuels Mtb’s metabolism. The mechanism that controls expression of the *mce1* loci in response to fatty acid depletion is unknown but we hypothesize that Mtb has an ability to sense metabolite pools to control expression of the Mce1 fatty acid transporter complex.

The Mtb cell envelope constitutes formidable barrier to the transport of any hydrophobic molecule. Actinomycetal bacteria with mycolic acid-containing cell walls that are capable of metabolizing cholesterol use the Mce4 complex to import the sterol (Mohn et al., 2008; Pandey and Sassetti, 2008; Perkowski et al., 2016) and this has lead to the idea that all four of the Mce complexes transport hydrophobic molecules across the Mtb cell wall. Various studies have linked Mce1 to Mtb virulence (Gioffre et al., 2005; Joshi et al., 2006; Shimono et al., 2003) but the function of the Mce1 complex was hitherto unknown. It has been reported that inactivating Mce1 in Mtb induces a lipid homeostasis defect and the accumulation of free mycolic acids in the Mtb cell wall (Cantrell et al., 2013; Forrellad et al., 2014). Based on these observations it was hypothesized that the Mce1 may transport fatty acids and/or mycolic acids across the cell wall/membrane of Mtb and it was reported that Mce1 mutant in Mtb displayed a minor defect in fatty acid uptake (Forrellad et al., 2014). We used assays comparable to those previously described and detected major perturbations in fatty acid assimilation in our Mce1 mutants (Figure 3 and Figure S11). We cannot fully explain the discrepancy between these findings but we have noticed that spontaneous mutants of Mtb unable to produce phthiocerol dimycocerosate (PDIM) assimilate less fatty acid compared to PDIM positive strains. Our work was conducted in a PDIM positive strain of Mtb Erdman, which unmasks defects associated with a Mce1 deletion. Studies with *Mycobacterium leprae* (Mlep) are consistent with the interpretation that Mce1 functions as a mycobacterial fatty acid transporter. The Mlep genome contains a single *mce* loci which is most similar to the *mce1* operon found in Mtb and Mlep is fully capable of importing and metabolizing palmitate following recoverey from animal tissues (Franzblau, 1988). The Mlep genome also encodes homolog of *lucA* (ml2337) suggesting that LucA is central for Mce1 function. The finding that LucA facilitates fatty acid and cholesterol import and stabilizes components of the Mce1 transporter provides additional evidence that Mce1 functions as a fatty acid transporter. Given that the Mce4 and Mce1 transporters demonstrates specificity for cholesterol and fatty acids, respectively (Figures 3E-H, S10, S11, S13) the fatty acid uptake defect in the Δ*lucA* mutant is likely a consequence of the degradation of Mce1 components (Figure 4D).

The *mce1-4* loci make up four separate operons in the Mtb genome and each operon encodes the putative protein subunits that likely comprise the individual Mce transporters. It is thought that each Mce transporter is substrate-specific and the individual subunits are predicted to perform discrete roles. Specifically, the *mce4* operon encodes two putative permease subunits (YrbE4A and YrbE4B), six cell wall associated Mce proteins (Mce4A-Mce4F), and two accessory subunits (Mam4A and Mam4B). We found that LucA interacts with Mam4B (Figure 4B and S6) and that Mam4B is required for the metabolism of cholesterol, however, the mutant lacking Mam4B is still capable of binding cholesterol (Figure S13). Based on this observation we propose that Mce4 imports cholesterol via a two-step process that involves cholesterol binding/shuttling across the cell wall followed by the final translocation through the cytoplasmic membrane delivering cholesterol into the cytosol (Figure S14). The accessory and permease subunits may participate in the final translocation of cholesterol across the cytoplasmic membrane while the Mce proteins likely participate in the binding/shuttling of cholesterol across the Mtb cell envelope. Binding/shuttling of lipids across the Mtb cell envelope may be analogous to what has been proposed for lipid trafficking by Mce proteins across the periplasm of gram-negative bacteria (Malinverni and Silhavy, 2009; Nakayama and Zhang-Akiyama, 2017; Thong et al., 2016). This two-step mechanism of nutrient uptake may be a generalizable mechanism for all Mce transporters. Additional support for the two-step model comes from our observation that the Mce proteins (Mce1A, Mce1D, and Mce1E) are degraded in the Δ*lucA* mutant and this strain is unable to bind/shuttle fatty acids across the Mtb cell envelope (Figure 4D). Lastly, we can detect a residual level of fatty acid and cholesterol uptake and metabolism in mutants lacking Mce1 and Mce4, respectively. It is very likely that compensatory systems function to import these lipids in the absence of Mce1 and Mce4 and we are currently testing this hypothesis.

Recently it was reported that orphaned mce associated protein Rv0199/OmamA facilitates cholesterol utilization in Mtb and stabilizes the Mce1 complex (Perkowski et al., 2016). We also found that LucA stabilizes subunits of the Mce transporters and interacts with Rv0199/OmamA, Rv0177/Mam1C and Rv3492/Mam4A (Figure 4B). This suggests that LucA is recruited to the Mce1 and Mce4 complexes via interactions with these accessory subunits to stabilize or assemble the transporters. Additionally, the putative ATPase, MceG/Rv0655 facilitates cholesterol uptake in Mtb (Pandey and Sassetti, 2008) and the stability of MceG increases when it is co-expressed with various YrbE permease proteins in Msm (Joshi et al., 2006). We found that the MceG protein is destabilized in the Δ*lucA* mutant, which could explain the cholesterol uptake defect in this mutant (Figure 4D). Homology searches based on 3-dimensional structures identified a putative protease inhibitor domain within the N-terminus of LucA (Kelley et al., 2015). We hypothesize that LucA may mediate local inactivation of a protease to maintain the integrity of the transporter complex. Regulating activity of the transport complexes through proteolysis would be a mechanism to rapidly halt nutrient uptake through the Mce transporters, however such a suggestion will require more research.

During growth in the presence of cholesterol Mtb shunts cholesterol-derived methylmalonyl-CoA (originating from propionyl-CoA) towards the increased synthesis of methyl-branched, cell wall polyketide lipids (Griffin et al., 2012; Jain et al., 2007; Yang et al., 2009). This metabolic shunting requires that sufficient fatty acid derived acyl-AMP primers are available to support biosynthesis of polyketide lipids (Quadri, 2014). When excess fatty acids are supplied to infected MΦs, Mtb can enhance the flux of propionyl-CoA into polyketide lipids such as PDIM during infection (Lee et al., 2013). It would be advantageous for Mtb to coordinate fatty acid assimilation to maintain the acyl-AMP pools required for efficient synthesis of methyl-branched lipids. Thus, coordination of cholesterol and fatty acid uptake by LucA could ensure that balanced levels of these nutrients are maintained for optimized metabolism.

The central carbon and lipid metabolic pathways of Mtb have emerged as potential drug targets (Rhee et al., 2011; VanderVen et al., 2015), therefore understanding the bottlenecks or weaknesses in these pathways will assist TB drug discovery. Additionally, the flux of fatty acids into TAG and central metabolism contributes to drug tolerance in Mtb (Baek et al., 2011), a phenotype that is further enhanced by immune pressure during *in vivo* infection (Liu et al., 2016). Targeting the specialized lipid metabolic pathways in Mtb that are involved in fatty acid and cholesterol utilization could be a viable strategy for the development of new drugs that reduce Mtb drug tolerance and augment current TB drug regimens. Our data have defined new participants in the complex processes of fatty acid and cholesterol assimilation by Mtb. A better understanding of the functional integration of Mtb’s specialized metabolic pathways is required to acquire a fuller appreciation of Mtb pathogenesis.

## Author Contributions

Conceptualization, E.V.N. and B.C.V.; Methodology, E.V.N., and B.C.V.; Formal Analysis, E.V.N. and B.C.V.; Investigation, E.V.N., C.R.M., T.L., N.S., W.L., S.C., K.M.W., B.C.V.; Writing – Original Draft, E.V.N. and B.C.V.; Writing – Review & Editing, E.V.N., B.C.V. and D.G.R.; Visualization, E.V.N. and B.C.V.; Funding Acquisition, D.G.R. and B.C.V.

## Acknowledgements

We thank Linda Bennett for excellent technical support, Robert Abramovitch, Shumin Tan, John Helmann, and Lu Huang for productive discussions, Maria Podinovskaia for assistance with imaging, Yancheng Liu for help with library construction for M-PFC. We also thank Adrie Steyn for the generous gift of the M-PFC vectors, Martin Pavelka for the aacC4 apramycin resistance cassette, and Christopher Sassetti for the Mce1 antibodies. This work was supported by the NIH grants (AI099569 and AI119122) to BCV and (AI080651) to DGR.

## Materials and Methods

### Bacteria and growth conditions

*M. tuberculosis* strains were routinely grown at 37°C in 7H9 (broth) or 7H11 (agar) media supplemented with OAD enrichment (oleate-albumin-dextrose-NaCl), 0.05% glycerol and 0.05% tyloxapol (broth). AD enrichment consisted of fatty acid free albumin-dextrose-NaCl. 7H9-based minimal medium is composed of Difco Middlebrook 7H9 powder 4.7 g/liter, 100 mM 2-(*N*-morpholino)ethanesulfonic acid pH 6.6, and carbon sources as indicated. Cholesterol was added to the liquid and solid media as tyloaxapol:ethanol micelles as described (Lee et al., 2013). Hygromycin 100 μg/ml, kanamycin 25 μg/ml, streptomycin 50 μg/ml, and apramycin 50 μg/ml were used for selection. For *E. coli* selection hygromycin was used at 150 μg/ml.

### Transposon screen and strain construction

A library of transposon mutants (∼10^5^) in a Δ*icl1* deficient strain of Mtb described by (Lee et al., 2013) was plated onto 7H11 OAD agar containing 100 μM cholesterol. Individual mutants were recovered in culture. Chromosomal DNA was isolated and the transposon insertion sites were PCR amplified and sequenced according to (Prod’hom et al., 1998). Mutant strains of Mtb were generated by allelic exchange (Mann et al., 2011) using a hygromycin resistance cassette mutant. Allelic exchange was confirmed by sequencing and/or Southern analysis using the Direct nucleic acid labeling and detection kit, GE Health Care. All the strains used in the study are summarized (Table S5).

### Radiorespirometry assays

Lipid oxidation was monitored by quantifying the release of ^14^CO_2_ from [4-^14^C]-cholesterol, [^14^C(U)]-palmitate, and [1-^14^C]-oleate by radiorespirometry as described (VanderVen et al., 2015). Briefly, Mtb cultures were pre-grown in 7H9 AD for 5 days, then incubated at OD_600_ of 0.7 in 5 ml 7H9 AD (albumin-dextrose-NaCl) medium supplemented with 1.0 μCi of radiolabeled substrates in vented standing T-25 tissue culture flasks. The culture flasks were placed in an air-tight vessel with an open vial containing 0.5 ml 1.0 M NaOH, sealed, and incubated at 37°C. After 5 hours, the NaOH vial was recovered, neutralized with 0.5 ml 1.0 M HCl, and the amount of base soluble Na_2_^14^CO_3_ was quantified by scintillation counting. The radioactive signal was normalized to the relative levels of bacterial growth by determining the OD_600_ for the bacterial cultures. % CO2 release was expressed as a ratio of normalized radioactive signal for each strain relative to the wild type control.

### Lipid uptake assays

Lipid uptake was quantified as described previously (Forrellad et al., 2014; Pandey and Sassetti, 2008) with slight modifications. Briefly, Mtb was cultured at an initial OD_600_ of 0.1 in 7H9 AD medium in vented standing T-75 tissue culture flasks. After 5 days, cultures were all normalized to OD_600_ of 0.7 in 8ml using spent medium, and 0.2 μCi of radiolabeled substrates was added to the cultures. After 5, 30, 60 and 120 min of incubation at 37°C 1.5 ml of the bacterial cultures were collected by centrifugation. Each bacterial pellet was washed thrice in 1 ml of ice-cold wash buffer (0.1% Fatty acid free-BSA and 0.1% Triton X-100 in PBS), fixed in 0.2 ml of 4% PFA for 1h. The total amount of radioactive label associated with the fixed pellet was quantified by scintillation counting. The radioactive signal was normalized to the relative levels of bacterial growth, i.e. to the OD_600_ of the bacterial cultures before addition of radioactive label. The uptake rate was calculated by applying linear regression to the normalized radioactive counts over time, and uptake efficiency was expressed as a ratio of uptake rate for each strain relative to the wild type control.

### MΦs isolation and culturing

MΦs were differentiated using bone marrow cells from BALB/c mice (Jackson Laboratories, USA) and maintained in DMEM supplemented with 10% heat inactivated fetal calf serum, 2.0 mM L-glutamine, 1.0 mM sodium pyruvate, 10% L-cell-conditioned media and antibiotics (100 U/ml penicillin and 100 mg/ml streptomycin) at 37°C and 7.0% CO_2_ for 10 days before infection. Human MΦs were differentiated from purified human peripheral blood mononuclear cells obtained from Elutriation Core Facility, University of Nebraska Medical Center and maintained in DMEM supplemented with 10% pooled heat inactivated human serum (SeraCare), 2.0 mM L-glutamine, 1.0 mM sodium pyruvate and antibiotics (100 U/ml penicillin and 100 mg/ml streptomycin) at 37°C and 7.0% CO_2_ for 7 days before infection. Media without antibiotics was used for infections with Mtb.

### Transcriptional profiling

Murine bone marrow-derived MΦs were seeded into two T-75 tissue culture flasks (1.5x10^7^ cells per flask) and infected with Mtb at a MOI of 4:1 for 3 days. Bacterial RNA was isolated, amplified, dye labeled, and hybridized to the microarray as described (Liu et al., 2016; Rohde et al., 2007). All transcriptional profile data has been deposited in the Gene Expression Omnibus database, accession number XXXX. The entire dataset is also available on ArrayExpress database (www.ebi.ac.uk/arrayexpress/), accession number XXXX.

### *prpD*’::GFP reporter assays

The promoter of *rv1130/prpD* was fused to GFP in a replicating vector that constitutively expresses mCherry (Table S7) To detect prpD promoter activity bacteria were grown in 7H9-based minimal medium containing 10 mM glucose for 5 days, washed twice with PBS 0.05% tyloxapol, and passed to medium containing 100 μM cholesterol or propionate at the indicated concentration for 24 hr. The bacteria were fixed with 4% paraformaldehyde (PFA) and GFP expression was quantified by flow cytometery on a BD Biosciences LSR II flow cytometer. To detect prpD promoter activity during infection bacteria grown 7H9-based minimal medium containing 10 mM glucose were used to infect murine MΦs at an MOI of 5:1. After 24 hr infection the MΦs were fixed with 4% PFA and scraped into 10 ml of PBS and suspended 1 ml of lysis buffer (0.1% SDS, 0.1 mg/ml Proteinase K in H_2_0). MΦs were lysed by 25 passages through a 25-gauge needle and the bacteria containing cell lysate was centrifuged and the pellet was retained and analyzed on a BD Biosciences LSR II flow cytometer. Flow cytometry data was analyzed using FlowJo (Tree Star, Inc).

### Imaging of intracellular lipid inclusions

Confluent monolayers of MΦs in Ibidi eight-well glass-bottom chambers were infected with bacteria at a MOI of 4:1. Extracellular bacteria were removed after 4 hours of infection. Infected MΦs were maintained in cell culture medium at 37°C and 7% CO_2_ for 3 days. Lipid inclusions of bacteria in MΦs were metabolically labeled with Bodipy-C16 (final concentration 20 μM) conjugated to 1.0% de-fatted bovine serum albumin (BSA) for a 30 minute pulse followed by a 1 hour chase with fresh media. Live-cell images were acquired as described (Podinovskaia et al., 2013). For lipid staining, MΦs were transferred onto sterile coverslips in 24-well plates, infected with Mtb for 3 days and fixed in 4% PFA followed by staining with BODIPY-493/503 (1.0 μg/ml, at room temperature for 1 hour). Post-acquisition, images were analyzed using Volocity (PerkinElmer Life Sciences).

### Flow cytometric quantification of assimilated lipids

Murine bone marrow-derived MΦs were seeded into T-150 tissue culture flasks (3×10^7^ cells per flask) and infected with Mtb at a MOI of 4:1. After 3 days of infection Bodipy-palmitate (final concentration 8 μM) conjugated to de-fatted 1% BSA was added to the cells for 1 hour pulse and then chased with cell media for another hour. The infected MΦs were scraped into 15 ml of homogenization buffer (250 mM sucrose, 0.5 mM EGTA, 20 mM HEPES, .05% gelatin, pH 7.0) and pelleted by centrifugation at 514xG (1,500 rpm, Beckman Allegra 6KR centrifuge, GH-3.8 rotor), followed by cell lysis by 70 passages through a 25 gauge needle. 5 ml of cell lysate was centrifuged at 146xG (800 rpm) for 10 minutes, supernatant (suspensions of phagosomes) was retained and treated with 0.1% Tween-80 at 4°C for 15 minutes to break-open Mtb containing vacuoles. Isolated bacteria were washed once in PBS+0.05% tyloxapol and fixed in 4% PFA. Flow cytometry data was collected on BD FACS LSR II and analyzed using FlowJo (Tree Star, Inc).

### Colocalization studies with Alexa647-dextran labeled lysosomes

At day 3 of infection, bone marrow derived MΦs were pulse labeled with 50 μg/ml Alexa647-dextran for 45 minutes and chased in fresh media for an additional 45 minutes. Following the chase period the infected cells were fixed and imaged by confocal microscopy. An extended focus merge of the two channels and the background was used to threshold the data as described (Costes et al., 2004) and colocalization was calculated using Volocity (PerkinElmer Life Sciences).

## Transmission electron microscopy

Imaging was conducted as described (Podinovskaia et al., 2013).

### Tetrahydrolipstatin treatment of infected MΦs

Infected bone marrow derived MΦs were treated with 100 μM tetrahydrolipostatin at 4 hours post infection and maintained in the medium throughout the infection. Infected cells were fixed at day 3, stained with Bodipy-493/503, and imaged by confocal microscopy.

### Protein fragment complementation screen

Library construction and the screen were performed as described (Singh et al., 2006). Briefly, Mtb genomic DNA was isolated and partially digested with AciI and HpaII, size fractionated (0.5-2 kb), and cloned into the ClaI site of pUAB300 upstream of the F_1,2_ domain of murine dihydrofolate reductase. *E*. coli MegaX^™^ DH10B^™^ T1 (Life Technologies) electrocompetent cells were used for transformation. In total 5×10^5^ independent clones were selected on LB hygromycin agar plates. The clone library was isolated by QIAGEN QIAfilter Plasmid Giga Kit and used to transform Msm mc^2^155 containing pUAB200 which co-expresses the bait protein LucA protein fused to the F3 (LucA-F_3_) domain of murine dihydrofolate reductase. In total 2×10^6^ clones were screened on plates containing trimethoprim 30 μg/ml. Clones containing fragments of the Mtb dihydrofolate reductase (*rv2763c/dfrA*) were identified by PCR and removed (85.8%). Inserts from the *dfrA*-negative clones were sequenced, and the only in-frame clone that was identified more than once (4 times) contained first 225 bp of *rv3492c/mam4B*.

### Spotting two-hybrid interaction assay

All Msm clones expressing the bait and prey constructs used for interaction assays are shown (Table S6). Msm was grown in modified 7H9-OAD media containing 2% glycerol, 0.5% additional glucose and 0.05% Tween-80 shaking at 37°C. Bacteria were diluted to ∼8.3×10^6^ bacteria/ml and grown 5 hours before diluting to ∼1×10^6^ bacteria/ml. Bacteria were diluted further to spot onto agar plates containing trimethoprim at 0, 15, and 30 μg/mL.

### Quantification of protein interactions

To quantify the strength of interactions we used an AlamarBlue based approach modified from (Singh et al., 2006). Briefly, log-phase cultures of Msm clones were transferred into 96-well microtiter plates at a density of 10^6^ of cells per well. Eight 2-fold serial dilutions of trimethoprim were made for each clone, from 600 to 4.69 μg/ml. The final volume in the wells was 200 μl. After 41 hours of incubation at 37°C, 30 μl 50% AlamarBlue (Life Technologies) (diluted with the media) was added to the wells and after incubating for 20 hours the fluorescence intensity was measured in Gemini EM Microplate Reader (Molecular Devices) with excitation at 530nm and emission at 590nm. 100% inhibition was assigned to the wells without cells, and 0% inhibition to the wells with cells without trimethoprim.

### Antibody generation and western analysis

Antibody for MceG was generated in rabbits using the peptide KAQAAILDDL conjugated to keyhole limpet hemocyanin by (Cocalico Biologicals). This peptide was used for antibody purification by immunoaffinity chromatography. For westerns bacteria were grown as for the lipid uptake assays. In the cases of chloramphenicol treatment, the antibiotic was added at 20 μg/ml 2 days prior to harvesting the bacteria. To generate lysates bacteria were washed twice with PBS 0.05% tyloxapol and fixed with 4% PFA for 1 hour. Fixed cells were washed twice in PBS 0.05% tyloxapol and lysed by sonication. Protein concentrations were determined by BCA (Thermo Fisher Scientific) and equivalent amounts of protein were resolved by SDS-PAGE and transferred to nitrocellulose membranes. The primary anti-Mce1A, anti-Mce1D and, anti-Mce1E antibodies were obtained from Christopher Sassetti (Feltcher et al., 2015), and the anti-GroEL *antibody was obtained from BEI resources*. A HRP-conjugated goat-anti rabbit IgG (Jackson ImmunoResearch) was used as the secondary antibody. ImageJ was used to quantify signals on Western blots.

### qPCR

Bacteria were cultured as described for western analysis and the RNA was extracted and analyzed as previously described (Abramovitch et al., 2011).

### Bacterial survival assay in MΦs

Confluent macrophage (human and murine) monolayers in 24-well dishes were infected with Mtb at a MOI 4:1 for murine cells and a MOI of 0.5:1 for the human cells. Extracellular bacteria were removed by washing with fresh media after 4 hours of infection. At indicated time points MΦs were lysed with 0.1% Tween-80 in water and the lysates were serially diluted in 0.05% Tween-80 in water. The lysates were plated on 7H11 OAD agar and CFU were quantified after 3-4 weeks incubation at 37°C.

### Mouse infections

Eight-week-old female C57BL/6J WT mice (Jackson Laboratories) were infected with 1,000 CFU of Mtb Erdman (wild type, Δ*lucA*, complement) via an intranasal delivery method as described (Sukumar et al., 2014). This was accomplished by lightly anesthetizing the mice with isoflurane and administering the bacteria in a 25 μl Volume onto both nares. At sacrifice, the lungs were removed and half of the lungs were fixed in 4% PFA overnight, while another half was used for bacterial load quantification. For the latter, lungs were homogenized in PBS 0.05% Tween-80 and plated on 7H11 OAD agar. CFU were quantified after 3-4 weeks incubation at 37°C.

### Lung histopathology

PFA fixed lung lobes were stained with hematoxylin and eosin by the Cornell Histology Laboratory. Stained sections were imaged using a Zeiss Axio Imager M1 equipped with an AxioCam Hrc camera.

### Ethics Statement

All animal care and experimental protocols were in accordance with the NIH “Guide for the Care and Use of the laboratory Animals” and were approved by the Institutional Animal Care and Use Committee of Cornell University (protocol number 2013-0030).

**Figure S1.**
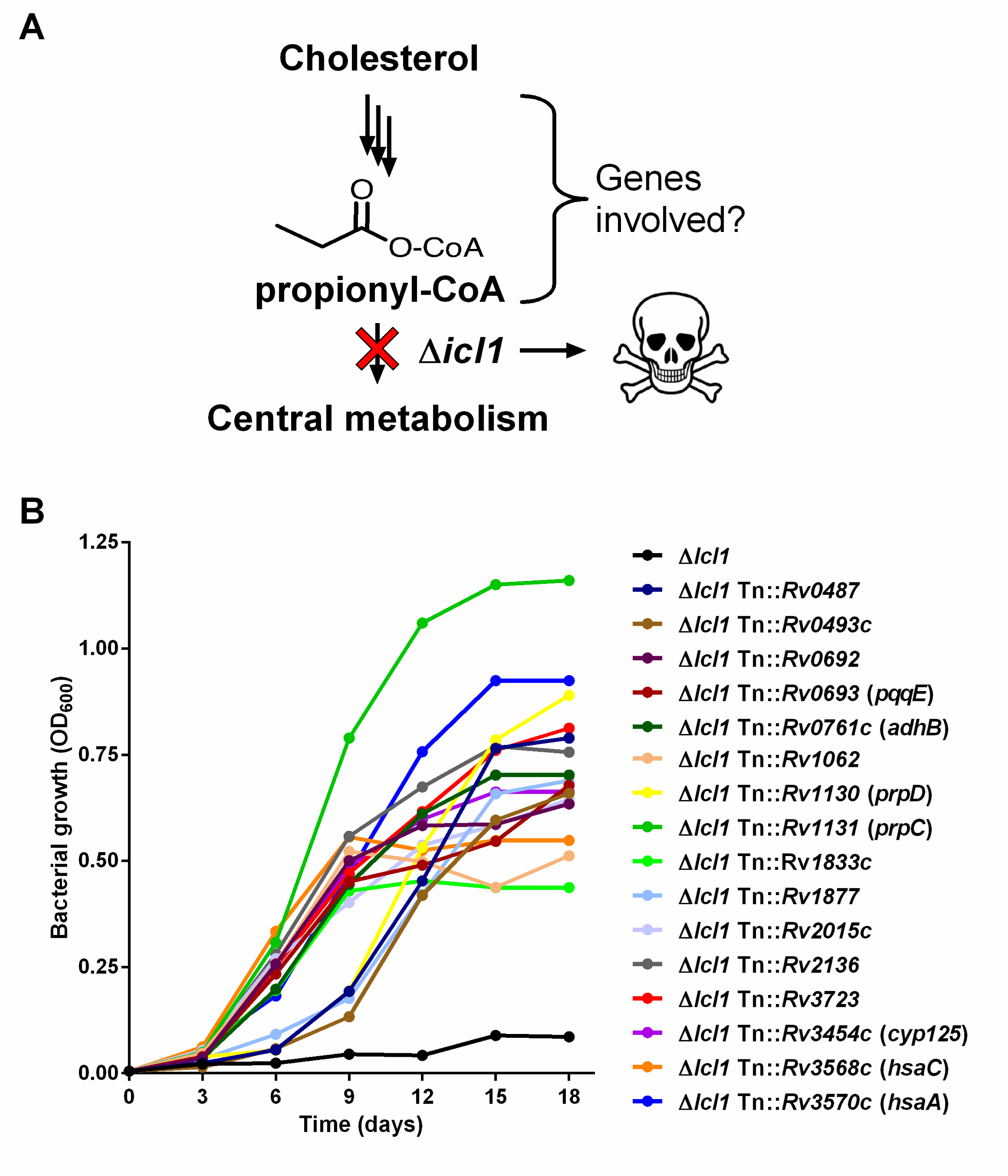
Mutations in cholesterol assimilation genes relieve cholesterol toxicity in Δ*icl1* Mtb. (A) Schematic detailing the source of cholesterol dependent toxicity in Δ*icl1* Mtb. (B) Transposon insertions rescue growth of Δ*icl1* Mtb. Representative clones carrying transposon insertions in the indicated genes in the Δ*icl1* background rescue bacterial growth in 7H9 OAD media supplemented with 100 μM cholesterol. Data are representative of two independent experiments.

**Figure S2.**
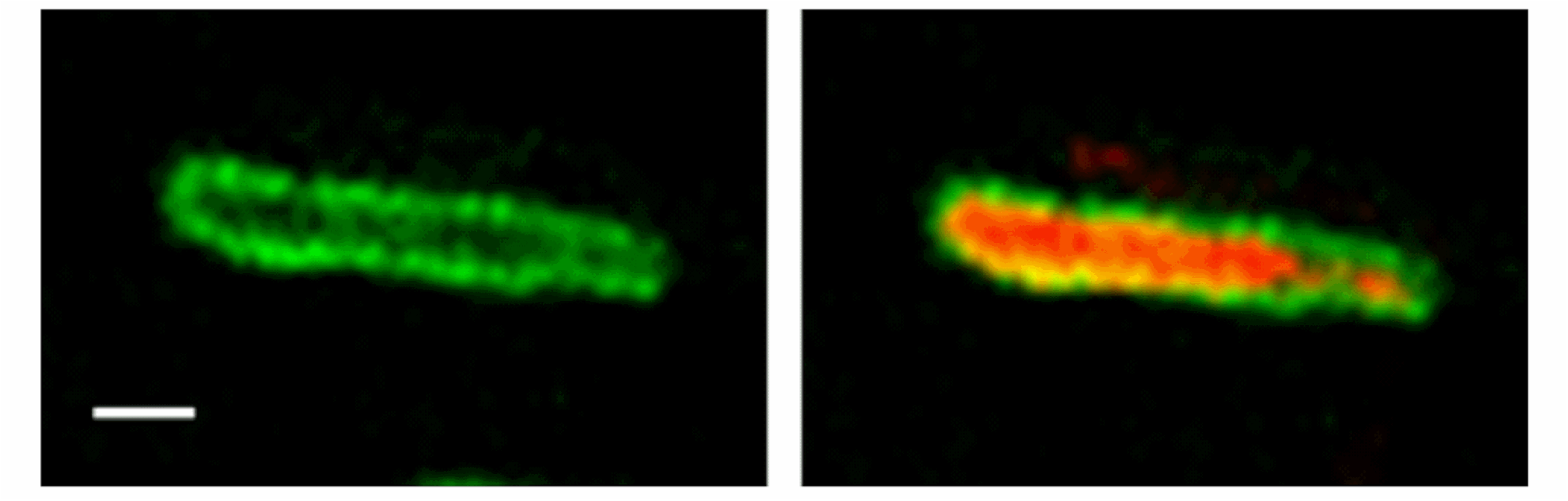
LucA-GFP has a cell surface localization pattern. Wild type Mtb constitutively expressing mCherry and LucA-GFP. Left panel a z-slice in the green channel alone (green = LucA-GFP) and the right panel overlaid green and red channels (red = mCherry M.tb) for the same optical slice. Scale bar 1.0 μm

**Figure S3.**
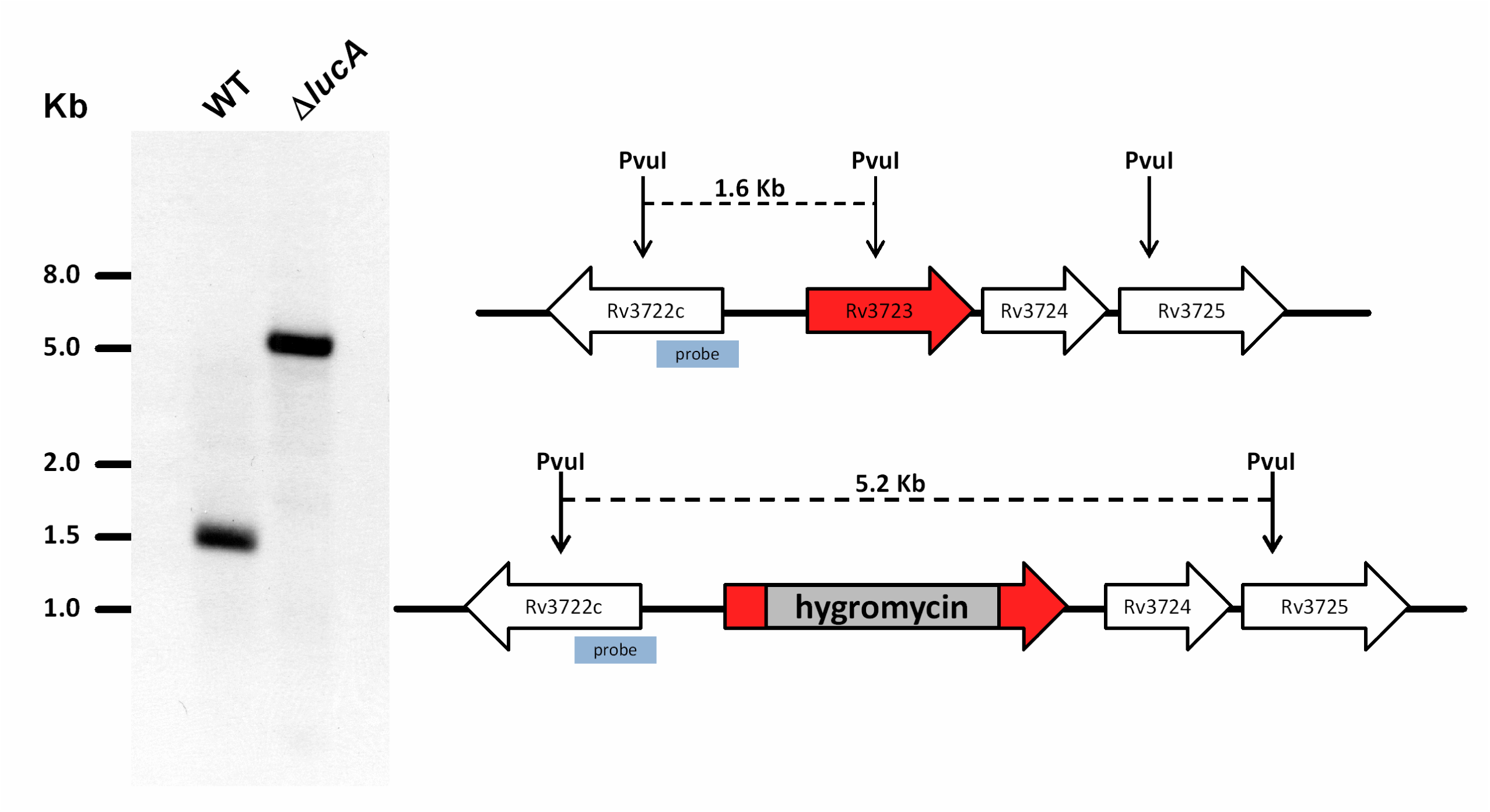
Confirmation of the ΔlucA mutant. Genomic DNA from Mtb was digested with PvuI and analyzed by Southern blotting using the indicated probe. Allelic exchange removes an internal fragment of *rv3723/lucA* containing a PvuI restriction site.

**Figure S4.**
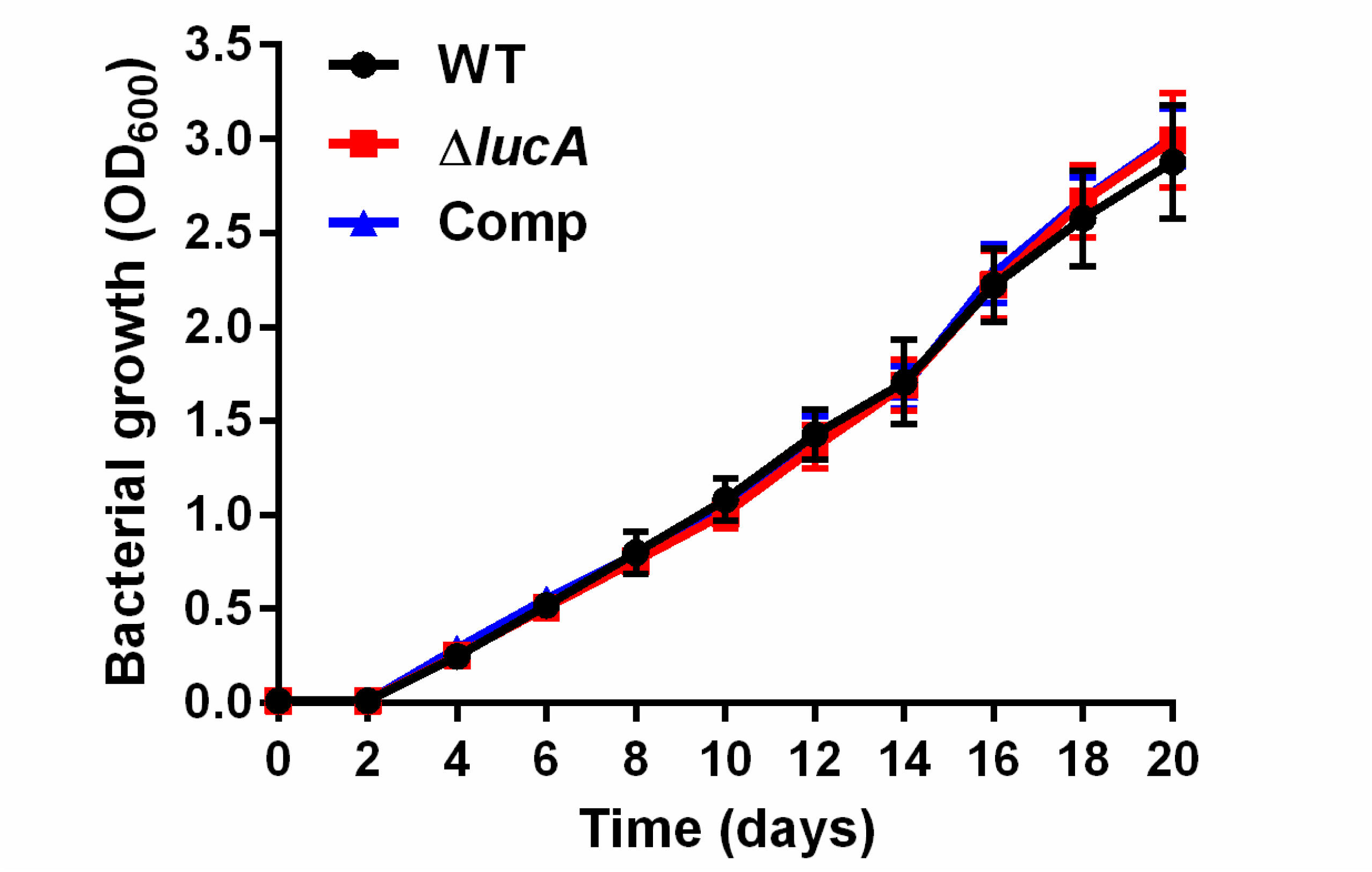
The Δ*lucA* mutant has no growth defect in 7H9 AD media. Bacterial growth was monitored across 20 days in 7H9 AD media. Data are means ± SD (n = 3)

**Figure S5.**
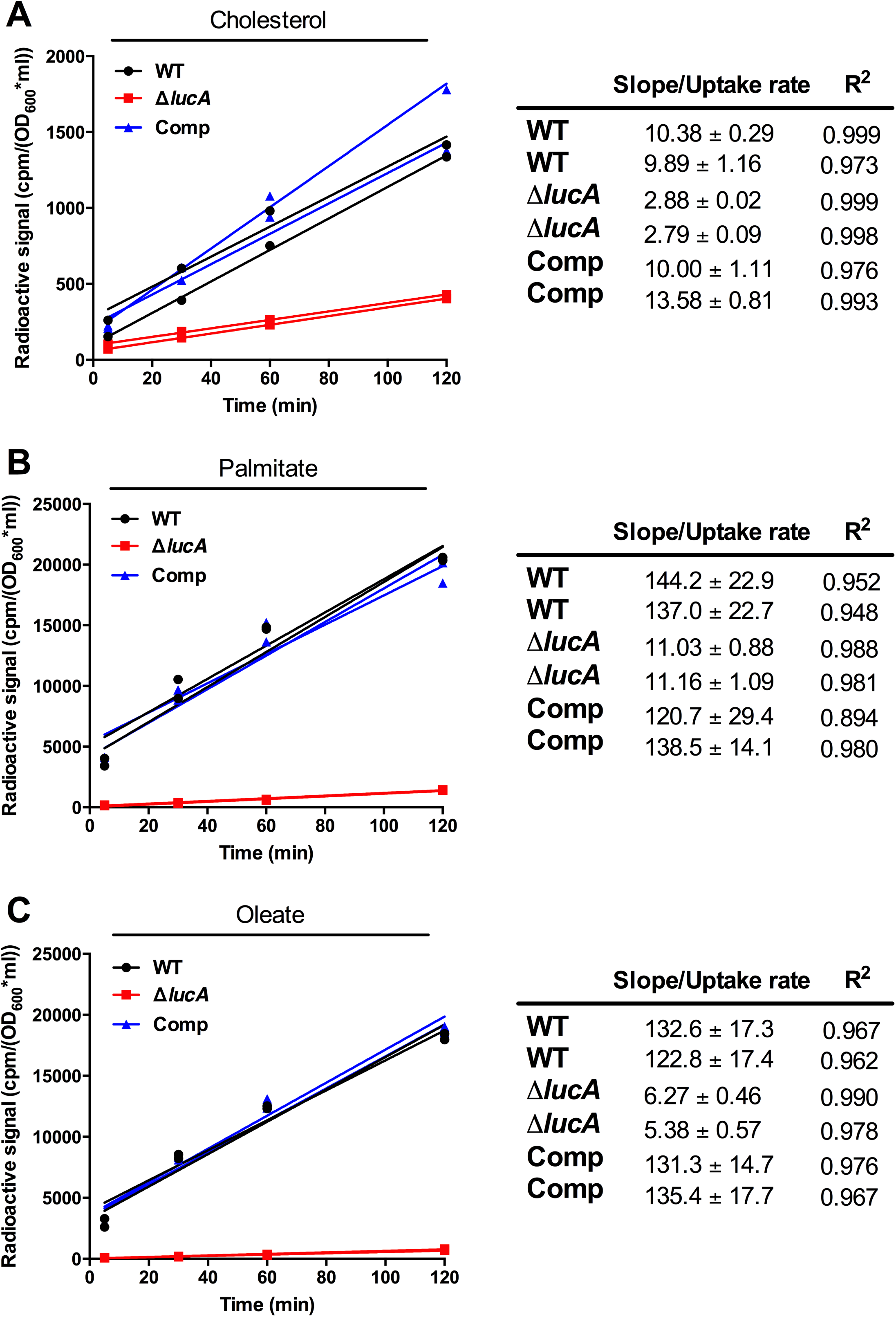
Raw radioactive counts and calculated lipid uptake rates. Raw counts from the cholesterol (A), palmitic acid (B), and oleic acid (C) uptake assays. Right, examples of how rate was calculated by applying linear regression to the cell-associated radioactivity counts. The slopes were used for quantification of the uptake efficiency. Data shown is representative of one independent experiment with two biological replicates. R^2^ indicates fit of linear regression.

**Figure S6.**
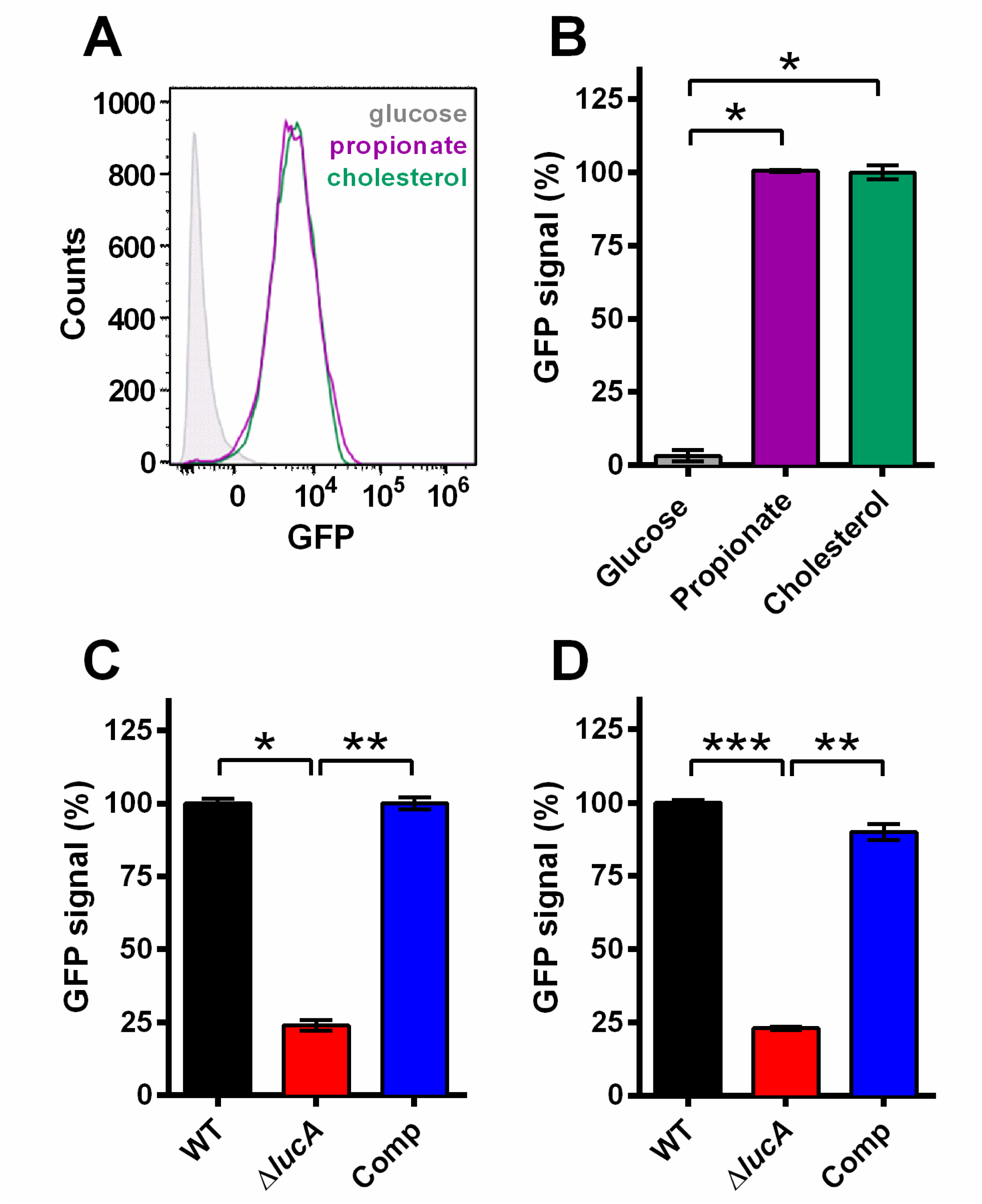
The Δ*lucA* mutant has defect in prpD gene expression. (A and B) Wild type Mtb carrying the *prpD* promoter reporter (*prpD*’::GFP *smyc*’::mCherry) expresses GFP in media containing cholesterol (100 μM) or propionate (300 μM). GFP signal was quantified by flow cytometery from 10,000 events. (B) The median GFP signal from wild type grown in cholesterol media was set to 100% and the GFP signals for bacteria grown in glucose and propionate was expressed as a ratio relative to the GFP signal from cholesterol media. (C) The Δ*lucA* mutant carrying the *prpD* promoter reporter has a defect in GFP expression when grown in cholesterol-containing media. WT was used as 100% to compare to the other strains. (D) The Δ*lucA* mutant carrying the *prpD* promoter reporter has a defect in GFP expression during infection of resting bone marrow derived MΦs. Bacteria expressing *PrpD*’::GFP were isolated from macrophages after day 1-post infection and analyzed by flow cytometry. WT was used as 100% to compare to the other strains. (B-D) Data are means ± SD (n=3). *p < 1*10^−6^, **p < 5*10^−6^, ***p < 5*10^−8^ (Student’s t test).

**Figure S7.**
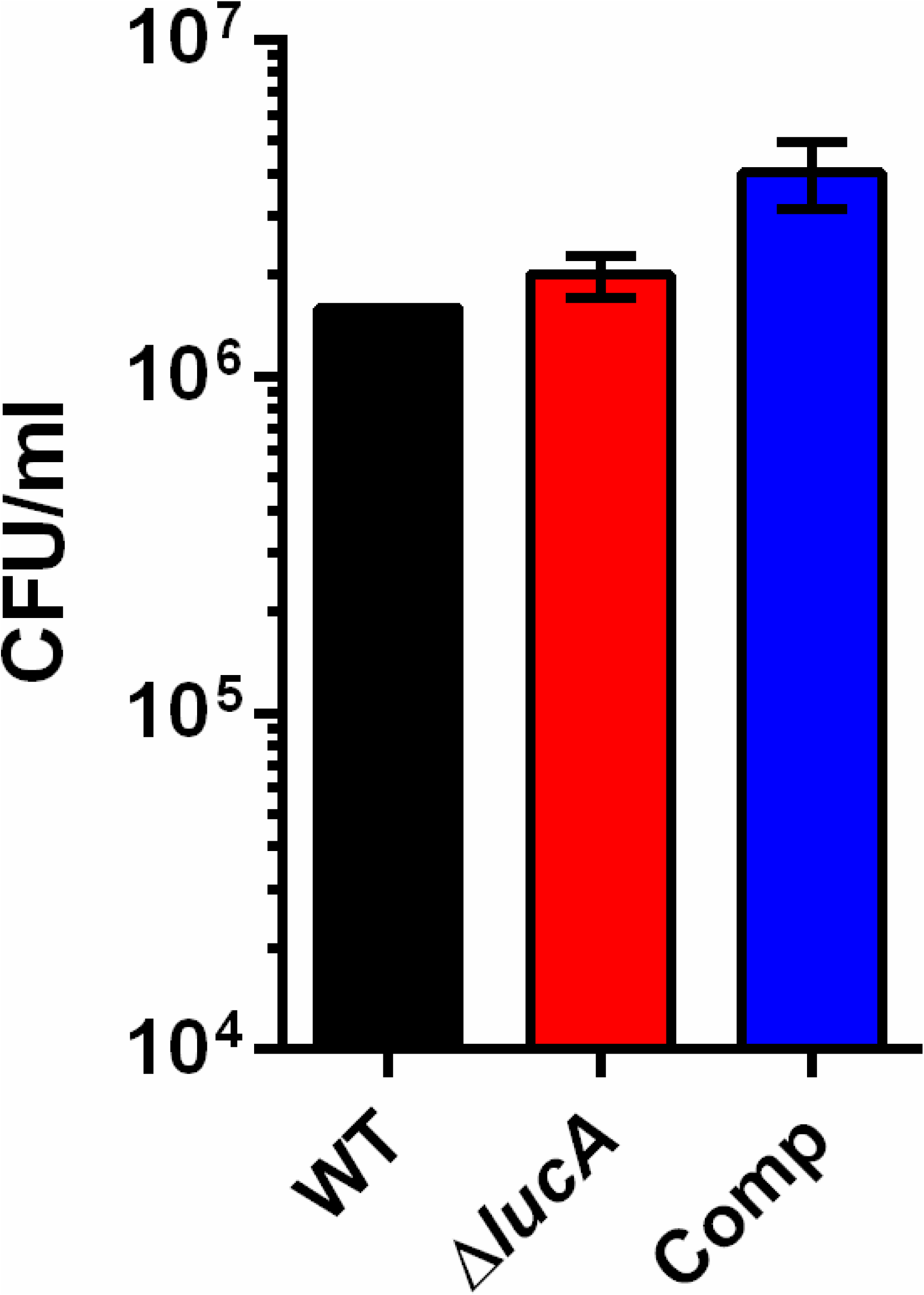
There is no defect in survival of Δ*lucA* mutant in MΦs at day 3 of infection. Bacterial viability determined at day 3-post infection from resting murine MΦs. Data are means ± SD (n = 3).

**Figure S8.**
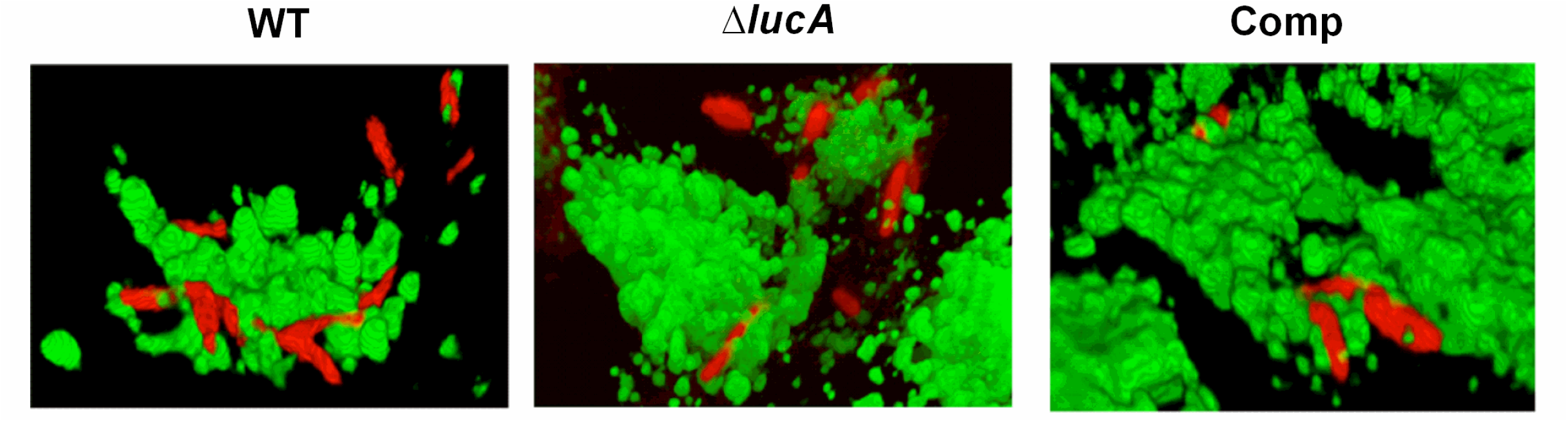
The Δ*lucA* mutant does not colocalize with pulse labeled lysosomes. Three-dimensional reconstructions of infected MΦs pulse-labeled with Alexa647-dextran (green = Alexa647-labeled) and (red = mCherry Mtb).

**Figure S9.**
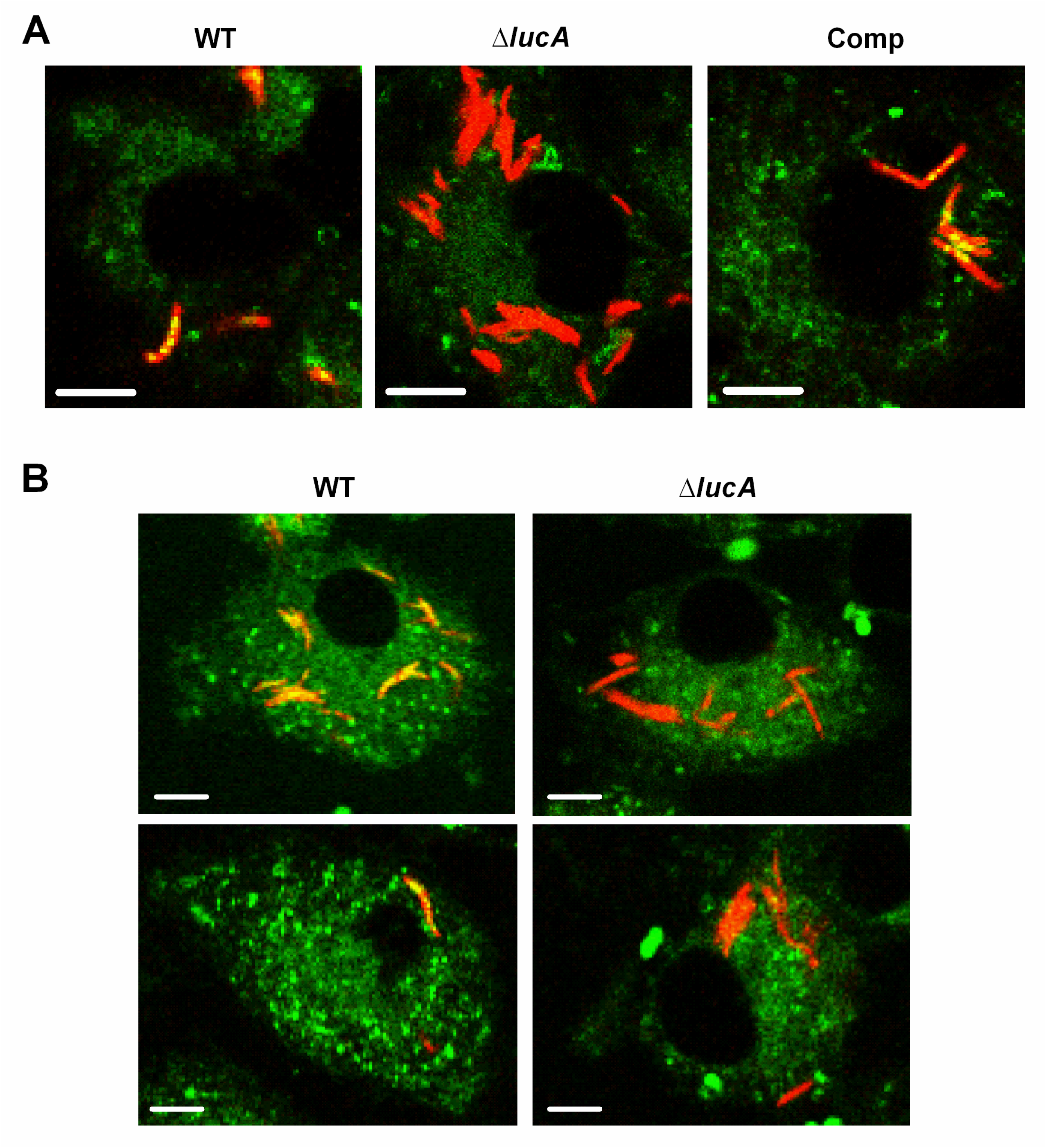
The Δ*lucA* mutant does not accumulate intracellular lipid inclusions that can be stained with Bodipy-493/503. (A) Infected bone marrow derived MΦs were fixed at day 3, stained with Bodipy-493/503, and imaged by confocal microscopy. Analysis of z-slices demonstrates that both wild type and the complemented mutant form intracellular lipid inclusions indicated by the punctate intracellular staining within the bacteria (red = mCherry Mtb) and (green = Bodipy-493/503), while Δ*lucA* mutant does not. Scale bar, 5.0 μm. (B) THL treatment does not alter levels of intracellular lipid inclusions in the Δ*lucA* mutant. Analysis of z-slices from treated cells demonstrates that wild type labels positive for intracellular lipid inclusions and the mutant does not. Lipid inclusions are identified by the punctate intracellular staining with Bodipy-493/503 within the bacteria (red = mCherry Mtb) and (green = Bodipy-493/503).

**Figure S10.**
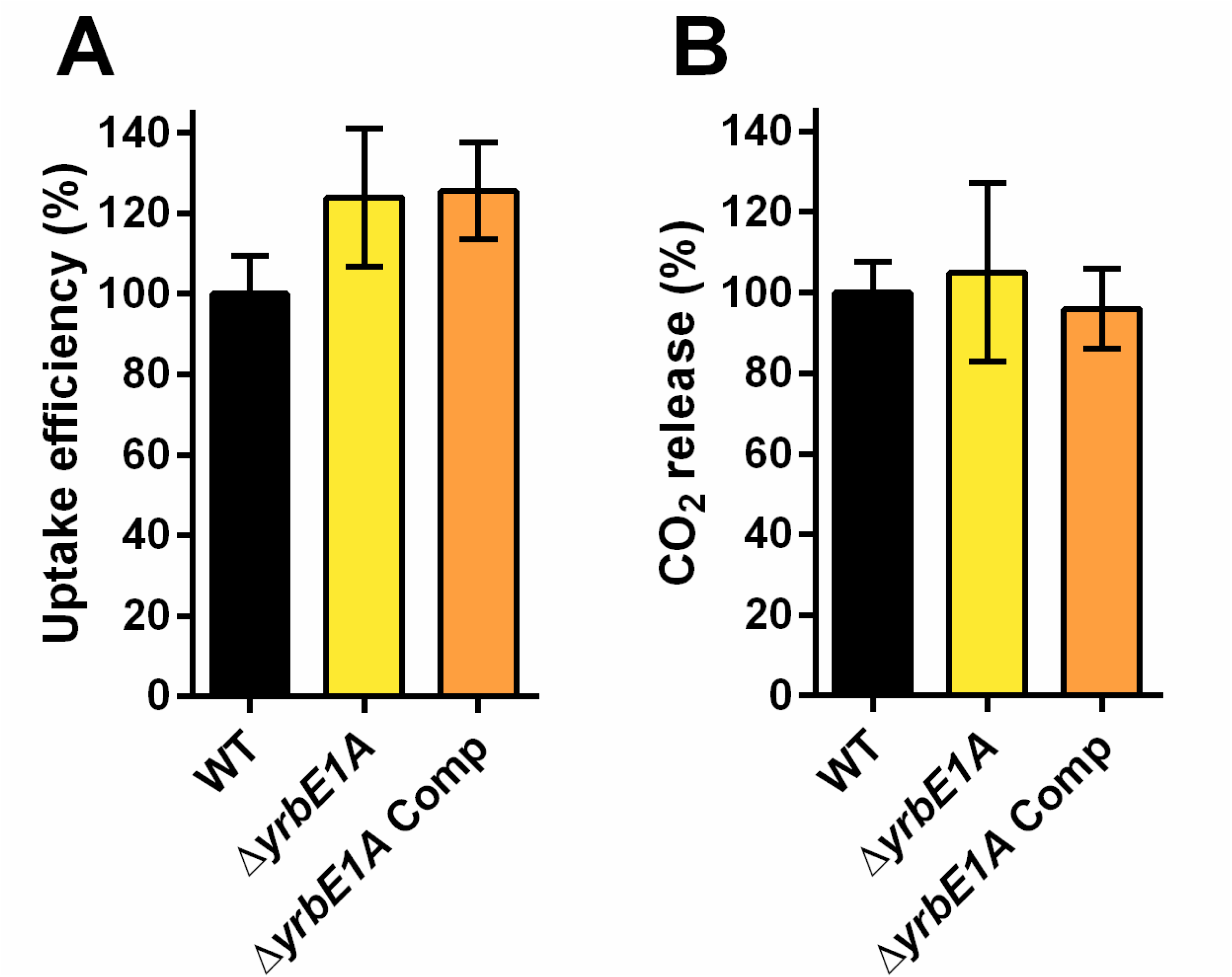
YrbE1A is not required for cholesterol assimilation in Mtb. (A and B) The Δ*yrbE1A* mutant has no defect in of cholesterol uptake (A) and metabolism(B). Data was calculated as in Figure 1B and 1C. Data are means ± SD (n ≥ 4).

**Figure S11.**
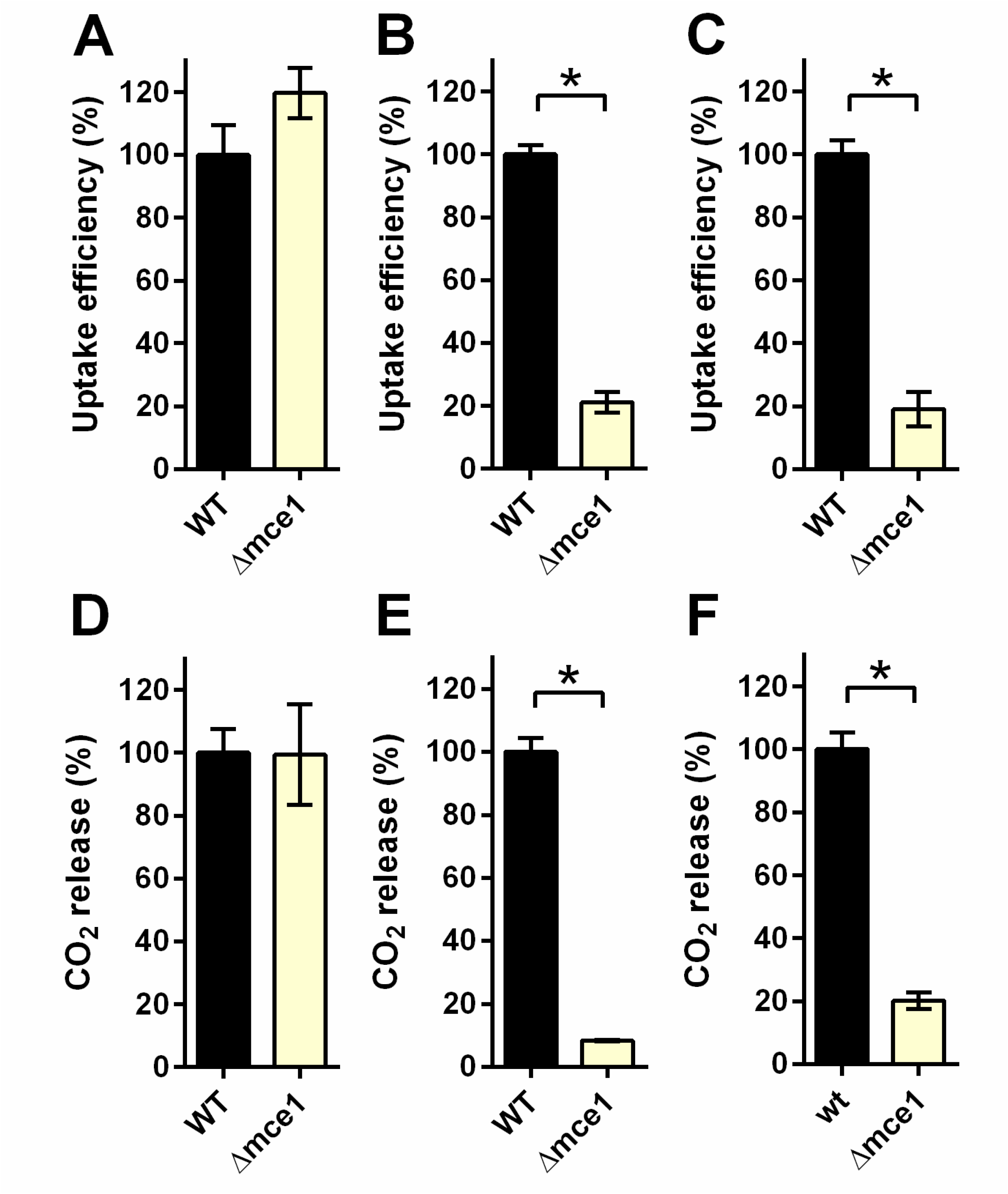
Deletion of the full Mce1 operon leads to fatty acid uptake defect. (A and D) The Δmce1 has no defect in cholesterol uptake (A) and metabolism (D). Data was calculated as in Figure 1B and 1C. (B and E) The Δmce1 is defective in oleic acid uptake (B) and metabolism (E). Data was calculated as in Figure 3E-H. (C and F) The Δmce1 is defective in palmitic acid uptake (C) and metabolism (F). Data was calculated as in Figure 3E-H. Data are means ± SD (n ≥ 4). *p < 0.0001 (Student’s t test).

**Figure S12.**
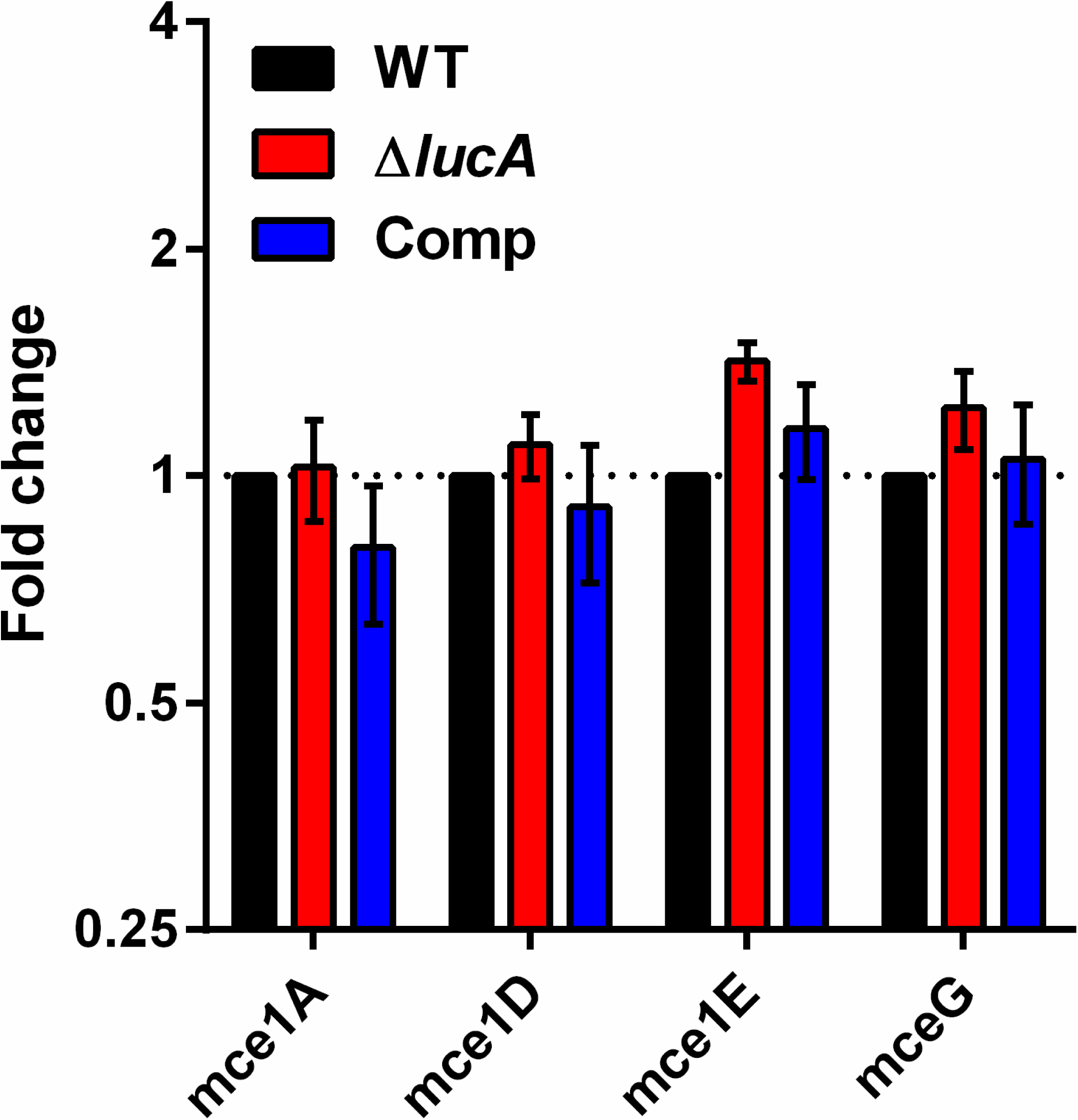
Deletion of LucA does not alter transcription levels of Mce1 and MceG. qPCR was performed using RNA isolated from Mtb. Fold change values were determined by normalizing transcript levels to the Mtb housekeeping gene, sigA. Data are means ± SD (n = 6)

**Figure S13.**
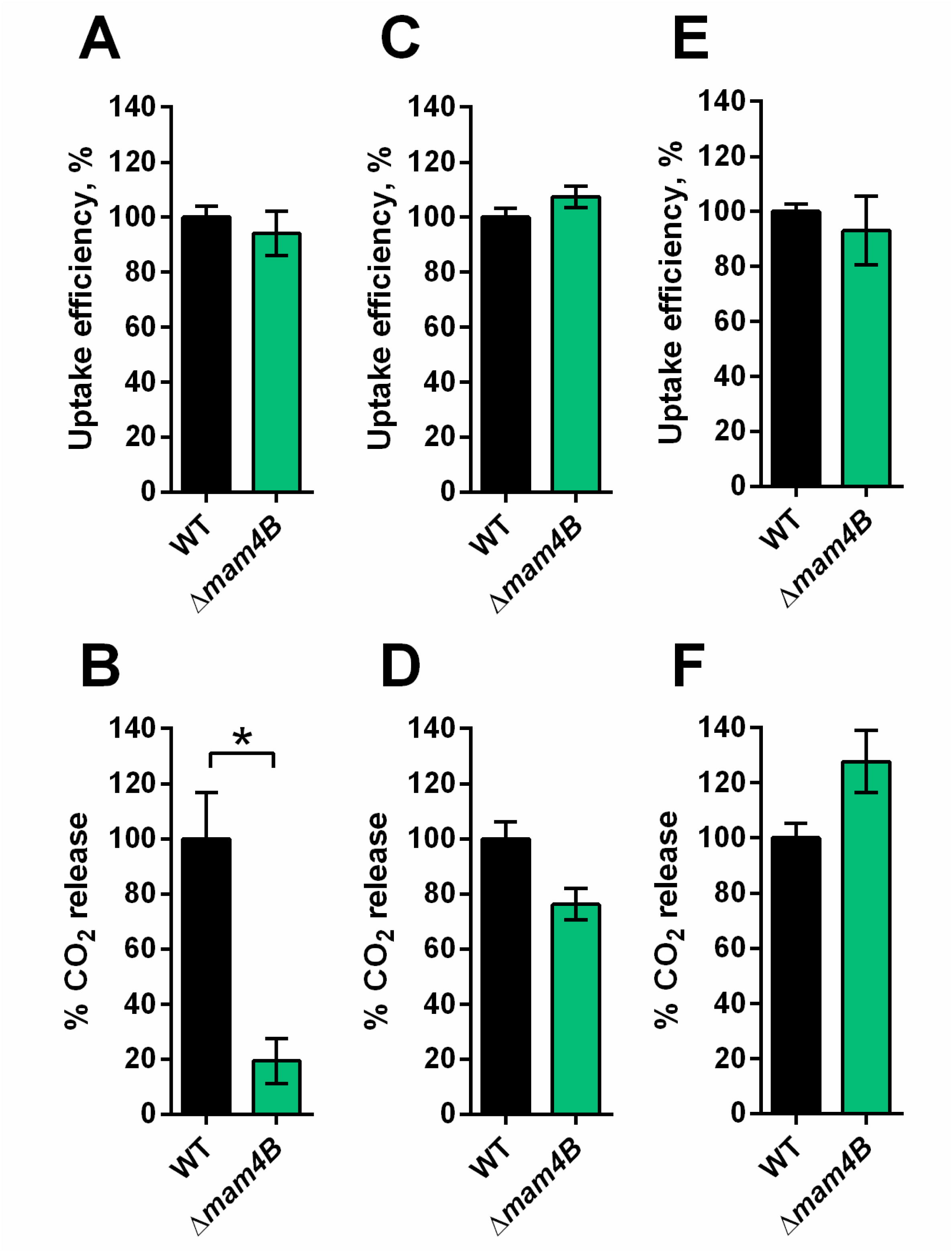
Mas4B is required for cholesterol assimilation. (A) The Δ*mas4B* mutant is capable of cholesterol uptake. Data are means ± SD (n = 4). Data was calculated as in in ure 1C. (B) The Δ*mas4B* mutant is defective in cholesterol metabolism. Data are means ± SD (n ≥ 4). Data was calculated as in Figure 1B and 1C. (C-D) The Δ*mas4B* mutant has no defect in oleic acid uptake (C) or metabolism (D). Data are means ± SD (n = 4). (E-F) The Δ*mas4B* mutant has no defect in palmitic acid uptake (E) or metabolism (F). Data are means ± SD (n = 4). (C-F) Data was calculated as in Figure 3E-H. *p < 0.0001 (Student’s t test).

**Figure S14.**
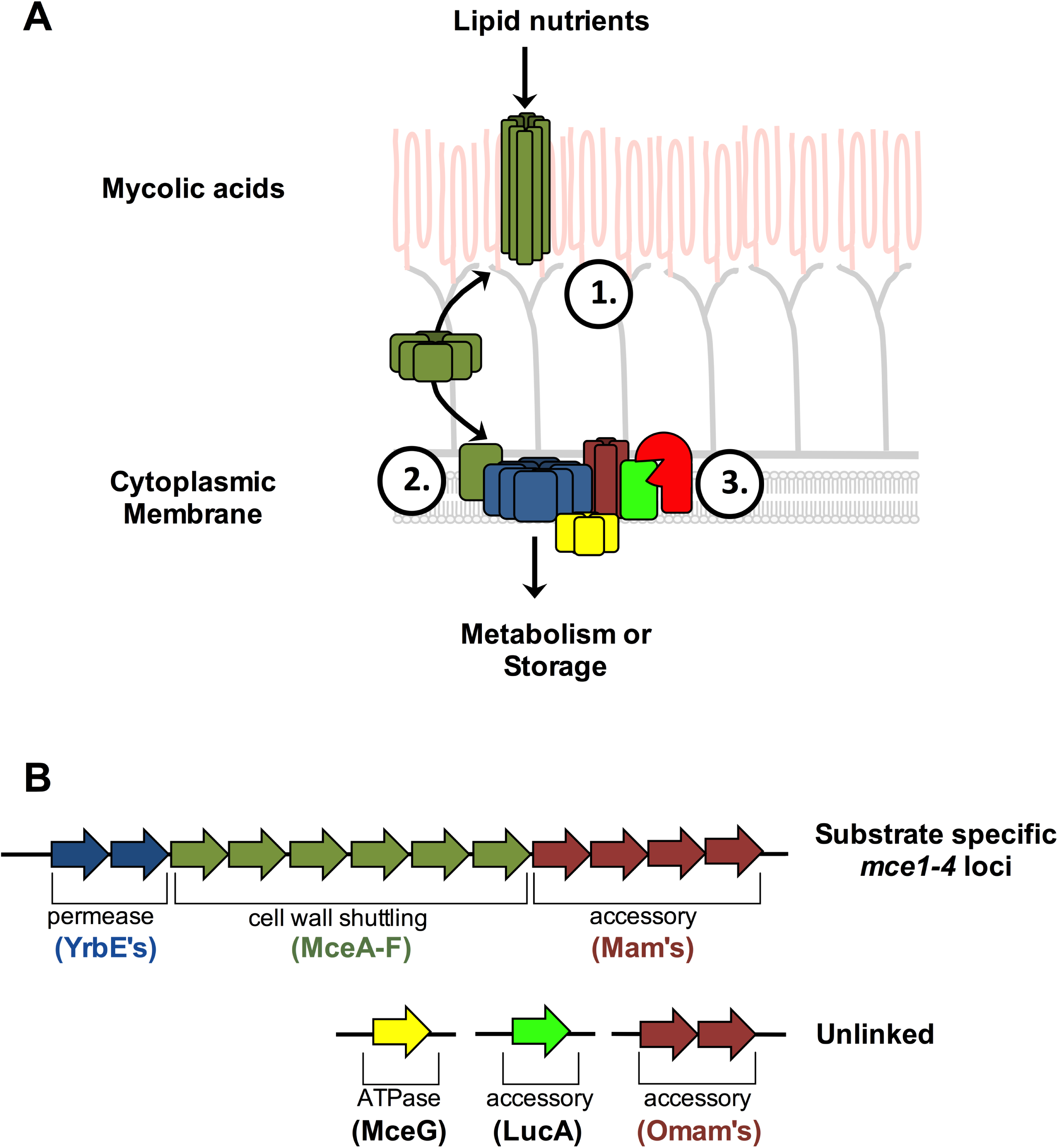
Hypothetical model for Mce-mediated transport of lipid nutrients into Mtb. (A) Proposed spatial arrangement and function for components of the Mce transporters. The secreted proteins (MceA-F) likely participate in the binding and shuttling of lipid nutrient across the mycolic acid layer and “pseudo-periplasmic space” of the Mtb cell wall, indicated as Step 1. These binding and shuttling events deliver the nutrient to specific cytoplasmic membrane proteins. The cytoplasmic membrane proteins function to translocate nutrients across the membrane indicated as Step 2. The permease subunits likely serve as the substrate pore and the accessory subunits likely act as an adapter to recruit additional proteins to the complex such as MceG and LucA which potentially regulate activity/stability of the transport complex. LucA likely interacts with and protects the Mce complexes from proteolytic degradation from an unknown protease indicated as Step 3. (B) Depiction of genome organization for known and predicted subunits involved in Mce1 and Mce4 mediated translocation of fatty acids and cholesterol respectively.

**Table S1. Mutants identified with the rescue screen in Δ*icl1* Mtb**

The disrupted gene is denoted along with the corresponding function for the mutated gene. The number of mutants indicates the total number of mutant clones identified and parentheses denote the number of independent insertions identified in the specific allele. Mutants indicated in red are predicted to be required for Mtb growth on cholesterol as a sole carbon source (Griffin et al., 2011).

**Table S2. Uptake rate calculations**

Cholesterol, palmitic acid or oleic acid uptake was quantified during incubation of Mtb with tracer levels of radiolabeled substrates for 2 hours. Linear regression was applied to the cell-associated radioactivity counts over time to quantify uptake rates. These uptake rates determined as the slopes were used for quantification of the uptake efficiency. Data is representative of one of the independent experiments with two biological replicates. R^2^ indicates fitness of linear regression.

**Table S3. Full bacterial transcriptional responses at day 3-post infection in murine bone marrow-derived MΦs**

**Table S4. Cholesterol-specific transcriptional response at day 3-post infection in murine bone marrow-derived MΦs**

**Table S5. Fatty acid-specific transcriptional response at day 3-post infection in murine bone marrow-derived MΦs**

**Table S6. Protein interactions constructs**

**Table S7. List of strains used in this study**

